# A lipid-structured model for macrophage populations in atherosclerotic plaques

**DOI:** 10.1101/557538

**Authors:** Hugh Z. Ford, Helen M. Byrne, Mary R. Myerscough

## Abstract

Atherosclerosis is a chronic inflammatory disease driven by the accumulation of pro-inflammatory, lipid-loaded macrophages at sites inside artery walls. These accumulations lead to the development of atherosclerotic plaques. The rupture of plaques that contain lipid-rich necrotic cores can trigger heart attacks and strokes via occlusion of blood vessels. We construct and analyse a system of partial integro-differential equations that model lipid accumulation by macrophages, including generating apoptotic cells and a necrotic core. The model includes the following cell behaviours: recruitment of macrophages into the plaque; macrophage ingestion of low density lipoproteins LDL and of apoptotic cells and necrotic material; lipid offloading to high density lipoproteins (HDL); macrophage emigration; and macrophage apoptosis and necrosis of apoptotic cells. With this model, we study how changes in parameters predict the characteristic features of plaque pathology. In particular, we find the qualitative form of lipid distribution across the macrophage population and show that high lipid loads can occur in the absence of LDL ingestion. We also demonstrate the importance of macrophage emigration in the model in mitigating and resolving inflammation and plaque lipid accumulation.

**Contributions:**

- HZF: conceptualisation, formal analysis, investigation, methodology, visualisation, writing— original draft preparation, writing—review and editing.
- HMB: conceptualisation, funding acquisition, methodology, project administration, resources, supervision, writing—review and editing.
- MRM: conceptualisation, funding acquisition, methodology, project administration, resources, supervision, writing—original draft, writing—review and editing.

## 1 Atherosclerosis and the role of lipid accumulation in macrophages

Atherosclerosis is a chronic inflammatory disease of the artery wall [1–4]. Inflammation occurs at sites inside the artery wall where modified low-density lipoproteins (LDL) accumulate after entering the wall from the bloodstream [5]. The immune response attracts circulating monocytes into such sites [6–8]. After crossing the endothelium (the layer of cells that line the lumen of the blood vessel) the monocytes enter the blood vessel wall and typically differentiate into macrophages that clear LDL via phagocytosis [9]. As the macrophages accumulate internalised lipid, they produce inflammatory signals that promote further monocyte recruitment [10, 11]. In this way, persistent LDL and monocyte influx into the artery wall produce large numbers of lipid-laden macrophages. This positive feedback creates a nonresolving inflammatory response [12] that thickens and expands the artery wall, to produce an atherosclerotic plaque. Plaques may continue to grow over the course of a lifetime and amass a build up of lipid-rich necrotic debris (called the necrotic core) via necrotic cell death of live and apoptotic macrophages [4, 13–15]. Plaques may rupture due to artery wall deterioration leading to the release of thrombotic (blood-clot forming) material. If these blood clots block arteries then they may cause heart attacks or strokes.

The intensity of inflammation, in general, depends on the number of LDL particles and macrophages inside the artery wall. The number of monocyte-derived macrophages inside the plaque (of the order of 10^6^ cells) is determined by the balance [9] between rapid monocyte recruitment (of the order of 10^5^ cells per day) [6–8], macrophage proliferation [16, 17], rapid programmed cell death (apoptosis) (of the order of a day to a week) [14] and/or emigration of macrophages out of the artery wall [18, 19]. Macrophages accumulate lipid via phagocytosis of both LDL and apoptotic cells. (Efferocytosis refers to phagocytosis of apoptotic cells.) LDL phagocytosis introduces lipid into the macrophage population whereas efferocytosis recycles and concentrates lipid that is already present in the macrophage population. Macrophages can offload cholesterol to high-density lipoproteins (HDL) [20, 21] and emigrating macrophages ferry their accumulated lipid out of the plaque [18]. Macrophages store accumulated internalised lipid in benign cytoplasmic lipid droplets (which gives them a “foamy” appearance) and will eventually accumulate cholesterol in the cell membranes and in the cytoplasm as crystals [22]. In this state, macrophages display elevated levels of inflammatory signalling, [23], apoptosis [24, 25] and may undergo necrotic cell death [26].

Most macrophages enter the plaque via monocyte recruitment and eventually undergo apoptosis. A study of myocardial infarction estimated that roughly 10% of macrophages emigrated from the inflamed tissue [8]. Thus, when cell numbers remain stable, apoptotic macrophages are produced almost as rapidly as monocytes enter the plaque. Rapid clearence of apoptotic cells (efferocytosis) prevents postapoptotic necrosis [14, 15, 27]; inefficient efferocytosis can lead to the accumulation of lipid-laden apoptotic macrophages and large amounts of cell necrosis [15]. Cell necrosis causes the release of lipid-rich cell contents into the extracellular space. This extracellular pool of thrombogenic and lipid-rich necrotic debris forms the necrotic core of the plaque. Large macrophage numbers and large necrotic cores are hallmarks of symptomatic plaques [4, 13].

There is increasing interest in mathematical modelling of the immunology of atherosclerotic plaques due to the dynamic nature of inflammatory macrophage populations [9]. Many mathematical models divide the macrophage population into “macrophage” and “foam cells”. This is true in ODE models [28], spatially resolved PDE models [29–31] and other computational models [32]. Yet it is clear that there is a continuous distribution of lipid loads across macrophage populations, stretching from monocytes with low lipid loadings and macrophages that can be labelled as foam cells in atherosclerotic plaques. The variation in lipid load across a population of macrophages suggests that it may be instructive to develop structured models in which the macrophages are characterised by their intracellular lipid load similar to adipocytes [33].

In this paper we develop and analyse such a mathematical model, explicitly accounting for lipid accumulation by monocyte-derived macrophages and its impact on the size and structure of the macrophage population and the generation of necrotic material in the plaque. At the heart of our model is a state variable which represents the quantity of lipid contained by monocyte-derived macrophages; this state variable plays a similar role to age in age-structured population models and size in size-structured models [34–37]. Applying principles of mass conservation yields a coupled system of time-dependent, non-local, partial integro-differential equations (PDE) that describe how the number of macrophages within the plaque and their lipid content change over time. The model resolves the dynamic interplay between monocyte recruitment, LDL phagocytosis, lipid accumulation by macrophages and apoptotic macrophages, macrophage apoptosis, macrophage emigration, efferocytosis of apoptotic cells, and necrosis of live and apoptotic macrophage. Our model predicts how the balance of these cellular processes influences the development of inflammatory states in atherosclerosis and, in particular, the accumulation of high numbers of macrophages with high intracellular lipid loads, and necrotic debris (Figure 1).

**Figure 1:**
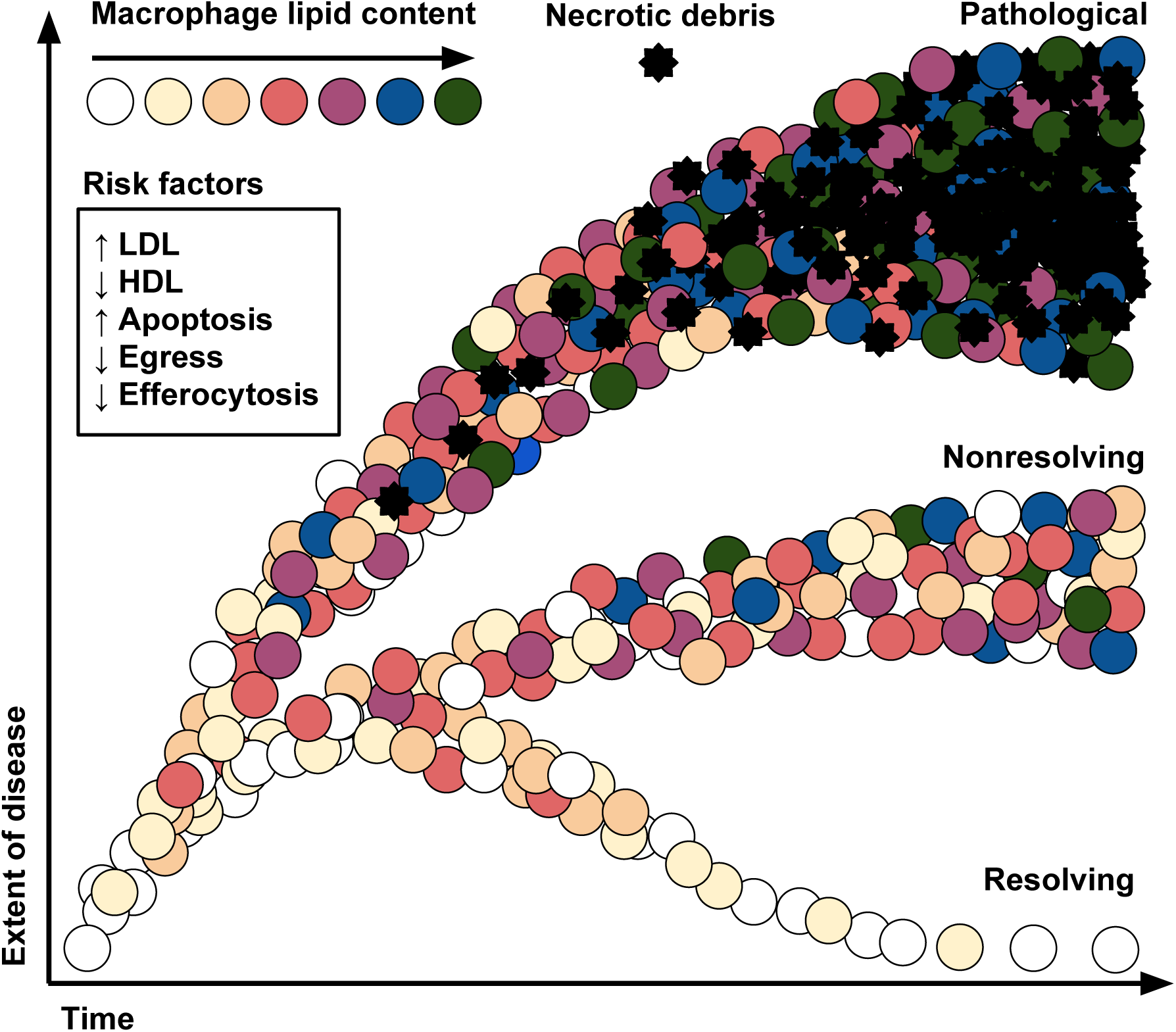
A schematic diagram showing how plaque composition changes over time as the severity of the disease increases. Monocyte-derived macrophages (circles) are allocated a shading to indicate their lipid content and necrotic material is indicated by black stars. Three distinct plaque fates are depicted: resolving plaques with low numbers of low lipid content macrophages and no necrotic debris; nonresolving plaques with a sustained number of macrophages with various lipid contents and no necrotic debris; and pathological plaques with elevated macrophage numbers shifted towards high lipid contents and a substantial necrotic core. Also indicated are the various risk factors are associated with the extent of the disease. These risk factors include elevated serum LDL levels and apoptosis rates and reduced serum HDL levels, cell emigration and efferocytosis.

The remainder of this paper is organised as follows. In section 2 we derive the general form of our model. It incorporates cellular processes which are known to play a significant role in the dynamics of lipid and macrophages, apoptotic cells and necrotic material within plaques. After nondimensionalising the governing equations, we present time-dependent numerical simulations. In section 3 we determine steady state solutions of the model and show how the distribution of internalised lipid across the macrophage population changes as the balance between macrophage immigration, efferocytosis, emigration, apoptosis and necrosis changes. In this way we reveal the subtle interplay between these processes and how their balance can be physiologically, pathologically or therapeutically altered to drive plaque pathogenesis or resolution. We also examine a special case for which there is no LDL (exogenous lipid) for macrophages to ingest but where efferocytosis of apoptotic cells cause macrophages to accumulate endogenous lipid derived from the membranes of macrophages that have since died and have been consumed. This special case could describe a situation where circulating monocytes are continually recruited to a tissue in response to foam cell that spontaneously form in inflammatory macrophage populations [10]. Finally, in section 4, we discuss our results and draw conclusions.

## 2 Model development

### 2.1 Model framework

We assume that the plaque is a dynamic mixture of live, apoptotic and necrotic macrophages and, for simplicity, spatial effects are neglected. Let *m*(*a, t*) *∈* R, *m*(*a, t*) *≥* 0 and *p*(*a, t*) *∈* R, *p*(*a, t*) *≥* 0 represent the number density in the plaque of live and apoptotic macrophages respectively with lipid content *a ∈* R, *a ≥* 0 at time *t ∈* R. Let *a*_0_ be the endogenous lipid content of macrophages so that *a ≥ a*_0_. That is, *a*_0_ represents the quantity of lipid in the cell’s membranes and other structures that is intrinsic to the cell itself and has not been accummulated by the macrophage ingesting other cell or lipids. Let *N* (*t*) *∈* ℝ, *N* (*t*) *≥* 0 represent the quantity of lipid in the necrotic core. Since necrotic material is acellular, *N*(*t*) is a function of *t* only. In constructing the model it will be useful to note that the total number of macrophages, *M*(*t*), and apoptotic cells, *P*(*t*), are given by:

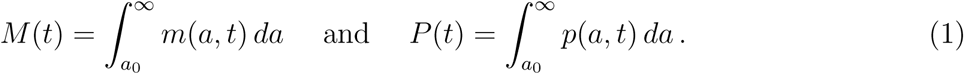

We will also use the total lipid load of all macrophages, *A_M_*(*t*), and of all apoptotic cells, *A_P_*(*t*), given by the first moments of *m*(*a, t*) and *p*(*a, t*) respectively:

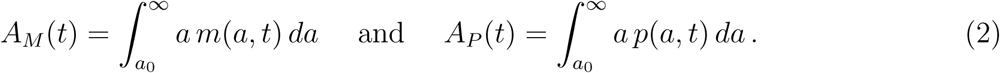

In this subsection we present the model equations in terms of the cellular processes that govern the time evolution of *m*(*a, t*), *p*(*a, t*) and *N*(*t*). The functional forms used to model these processes are outlined in subsequent subsections.

Our mathematical model comprises a system of two partial differential equations and one ordinary differential equation for *m*(*a, t*), *p*(*a, t*) and *N*(*t*) respectively:

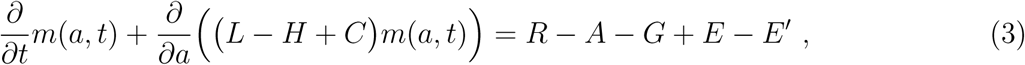

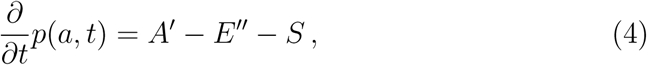

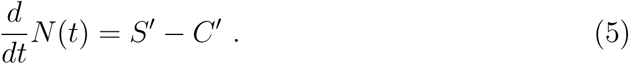

In equations (3)-(5), *A*, *A*′, *C*′, *E*, *E*′, *E*′′, *G*, *R*, *S* and *S*′ represent source and sink terms that drive changes in the distributions of live and apoptotic macrophages and the amount of necrotic material within the plaque.

The terms appearing in the advection term on the left hand side of equation (3) model processes which alter the lipid load of macrophages incrementally due to uptake of LDL (at rate *L*), offloading intracellular lipid to HDL (at rate *H*) and lipid uptake due to phagocytosis of necrotic material (at rate *C*).

Macrophage lipid loads also change when a macrophage ingests an apoptotic cell (efferocytosis). In this model, we assume that macrophages ingest whole apoptotic cells and acquire the entire lipid content of the apoptotic cell. In mathematical terms, efferocytosis produces non-local change as the lipid content, *a*, of the consumer cell increases instantaneously by a finite amount equivalent to the lipid content of the consumed apoptotic cell. In equation (3) *E* represents the rate at which macrophages of lipid load *a* are created through efferocytosis while *E*′ represents the rate at which macrophages of lipid load *a* move to a higher load through efferocytosis. The term *E*′′ in equation (4) represents the rate at which apoptotic cells with lipid content *a* are removed via efferocytosis.

In equation (3), macrophages are assumed to enter the plaque by crossing the endothelium and arriving in the artery wall at a rate *R*. We also assume that macrophages emigrate out of the artery wall at rate *G* and undergo apoptosis at rate *A*. We remark that the sink of live macrophages due to apoptosis in equation (3) is balanced by the source term *A′* in equation (4) for apoptotic cells.

Apoptotic cells that are not ingested by macrophages become necrotic at rate *S* and act as a source of lipid flowing into the necrotic core at rate *S*′. Material in the necrotic core is ingested and cleared by macrophages at rate *C*′.

In the next section, we introduce functional forms for each of these rates.

### 2.2 Specifying the functional forms for the model processes

#### 2.2.1 Lipid flux into and out of macrophages via LDL and HDL (***L*** and ***H***)

We suppose that serum entering into the artery wall at a fixed rate *σ* (unit volume per unit time) contains fixed concentrations of LDL and HDL which we denote by 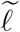 and 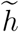 (particles per unit volume) respectively. We further assume that LDL particles contain a fixed amount of lipid *l*_0_ (lipid molecules per LDL particle), that they rapidly become modified once inside the artery wall (e.g. by oxidation) and are consumed by macrophages at a constant rate *ω* (per cell per unit time). Let *l*(*t*) be the number of LDL particles in the tissue at time *t*. If we assume that macrophages consume LDL at a rate proportional to the availability of LDL and independent of macrophage lipid load, then the change in *l*(*t*) is given by

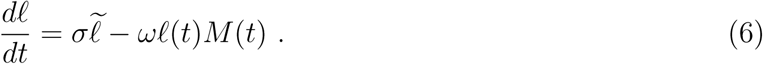

Rates of LDL ingestion occur on the scale of minutes-to-hours whereas rates of macrophage apoptosis occur on the scale of days-to-weeks [8, 38]. We assume that changes in *l* occur over a much shorter timescale than the timescale of the cell dynamics such that changes in cell numbers *M* produce rapid changes in LDL particle numbers *l*. Of interest here is the longer timescale associated with the cell dynamics; on this long timescale, the LDL dynamics are fast and we may make a quasi-steady state approximation. Setting 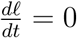 in equation (6) we deduce that:

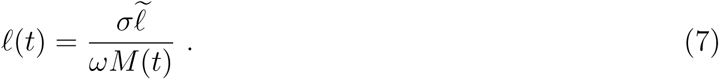

If we assume that the rate at which macrophages ingest lipid from LDL is proportional to the number of LDL particles in the tissue and the lipid load of each particle, then, from equation (7), we have that the total rate of lipid influx into each macrophage via LDL ingestion in equation (3) is:

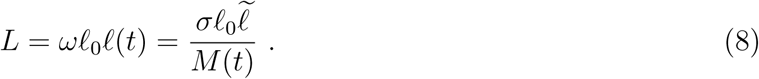

Once inside the artery wall, HDL particles take up lipid (principally cholesterol) from macrophages. We assume that macrophages offload lipid to HDL particles at a fixed rate *ξ* (lipid molecules per cell per HDL particle per unit time). If *h*_0_ (lipid molecules per HDL particle) is the lipid capacity of an HDL particle then the time taken for a HDL particle to be filled with lipid is *h*_0_*/*(*ξM*(*t*)) (unit time); assuming all macrophages have accumulated lipid to offload. If we assume that HDL particles become fully loaded with lipid before leaving the artery wall then the number of HDL particles *h*(*t*) inside the artery wall is modelled by:

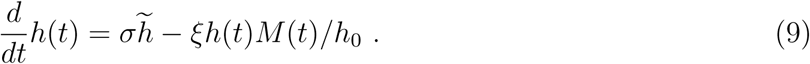

We assume that the LDL and HDL dynamics have a similar short timescale [38] such that the HDL dynamics are fast and we may make a quasi-steady state approximation. Setting 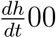 in equation (9) we deduce that:

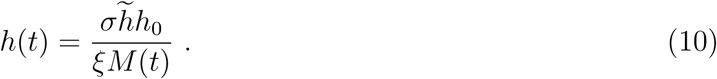

As we assume that individual macrophages offload lipid to HDL at a rate proportional to the number of HDL particles in the tissue, from equation (10), we have that the total rate of lipid efflux from each macropahge via HDL in equation (3) is:

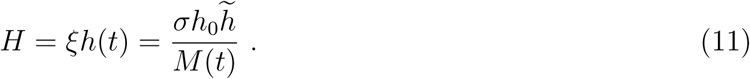

Combining (8) and (11) we deduce that the net lipid influx into macrophages is given by:

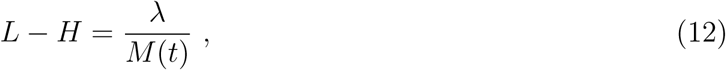

where 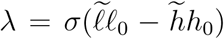 is the net flux of lipid into the macrophage population. The sign and magnitude of *λ* is determined by the relative serum concentrations of LDL and HDL particles 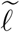 and 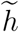 respectively and the LDL lipid content *l*_0_ and the lipid capacity of HDL *h*_0_. We note that the net rate at which macrophages take up lipid from lipoproteins depends on the total number of macrophages in the tissue.

#### 2.2.2 Monocyte recruitment from the vasculature

The recruitment of monocytes across the endothelium and into the artery wall is driven by cytokines such as interleukin 1 (IL-1) and monocyte chemoattractant protein 1 (MCP-1), that are produced by macrophages in response to lipid ingestion or high lipid loads [39, 40]. Once in the plaque, monocytes differentiate into macrophages.

We assume in the following derivation that there is only one generic cytokine, concentration *c*. We neglect cytokine production from other sources such as T-cells and necrotic or apoptotic cells. We assume that this cytokine is produced at rate *f* (*a*) (per cell per unit time) by macrophages with lipid load *a*. In general, *f* (*a*) is an increasing function of *a*. We also suppose that the cytokine degrades and/or leaves the system at rate *ζ* (per unit time). The time evolution of *c*(*t*) can therefore be modelled by the following integro-DE:

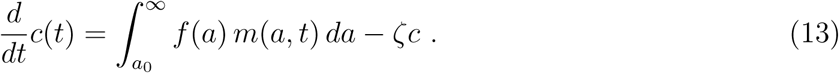

If the timescales on which cytokines are produced and degraded are much shorter than the timescale on which macrophage levels change, then as before we may assume that 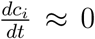 in which case equation (13) gives:

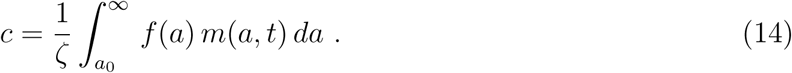

We assume that the rate of monocyte influx into the artery wall (that is, the recruitment rate) is a saturating function of *c*. Let *α* (cells per unit time) be the maximum recruitment rate and *K* be the cytokine level at which the recruitment rate is half maximal. Then the rate of monocyte recruitment is given by:

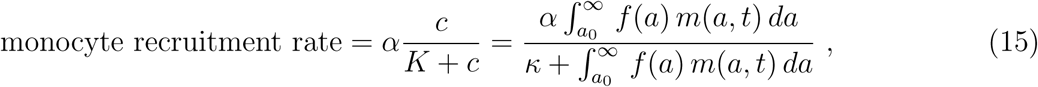

where *κ* = *Kζ*. For *n* cytokines with concentrations *c_i_*, 1 *≤ i ≤ n*, a similar analysis produces a similar recruitment function with the interpretation of *κ* and *f* (*a*) suitably adjusted.

For simplicity, we now assume that macrophages produce cytokines at a rate proportional to their accumulated lipid content so that *f* (*a*) = *a − a*_0_. If we assume further that monocytes enter the system with a fixed amount of endogenous lipid *a* = *a*_0_ then using equation (15) we arrive at the following functional form for *R*, the rate of recruitment of monocytes in equation (3):

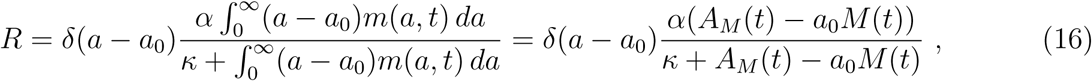

where *δ* is the Dirac delta function and *M*(*t*) and *A_M_*(*t*) are defined in equations (1) and (2). We remark that this source term can alternatively be cast as a boundary condition at *a* = *a*_0_. In all subsequent analysis, monocyte recruitment will be incorporated via a boundary condition at *a* = *a*_0_.

#### 2.2.3 Macrophage apoptosis (*A* and *A*′)

We suppose that macrophages undergo apoptosis at a constant rate *β* (per unit time) and produce equal numbers of apoptotic cells with equal lipid content so that:

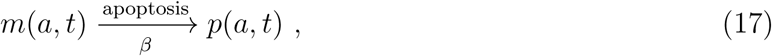

and the functional forms for *A* and *A*′ in equation (3) and (4) are:

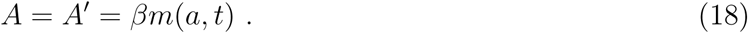

#### 2.2.4 Emigration of macrophages from the artery wall (*G*)

We assume that live macrophages emigrate from the artery wall at rate *γ* (per unit time) independently of their lipid content. In this case, the emigration sink term *G* in equation (3) is:

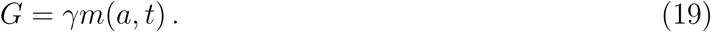

We emphasise that migrating cells remove cellular lipid from the artery wall and, as a result, emigration plays a key role in determining the overall lipid burden in the plaque and, in particular, the size of the necrotic core.

#### 2.2.5 Efferocytosis (*E*, *E* ′ and *E*′′)

We assume that apoptotic macrophages are typically consumed whole by live macrophage via efferocytosis. Efferocytosis transfers the lipid contents of an apoptotic cell, including its endogenous lipid, to the live macrophage that consumes it. We suppose that live macrophages consume apoptotic cells at a constant rate *η*, independent of the lipid contents of the apoptotic and live cells. We assume further that, when live macrophages with lipid content *a* consume apoptotic cells with lipid content *a*′, a more heavily loaded live macrophage with lipid content *a* + *a*′ is produced. Thus, we have:

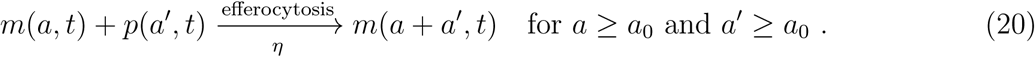

Since macrophages ingest apoptotic cells with lipid load *a*′ ∈ (*a*_0_, *∞*) the rate *E*′ at which live macrophages with lipid load *a* are lost due to efferocytosis is:

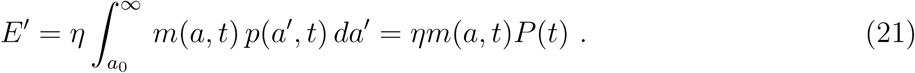

Macrophages that are removed from the population with lipid load *a* following efferocytosis produce macrophages with higher lipid loads. Thus the efferocytosis source term *E* in equation (3) is modelled by a convolution between the number of live *m*(*a, t*) and apoptotic *p*(*a, t*) macrophages with lipid content *a*′ and *a − a*′ respectively with *a′* < *a* − *a*_0_:

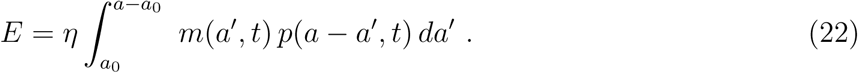

The rate *E*′′ at which apoptotic cells with lipid load *a* are removed as a result of efferocytosis is defined in a similar way so that:

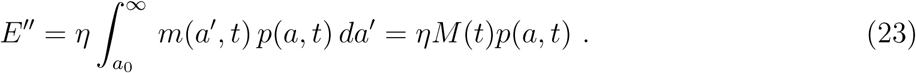

#### 2.2.6 Necrosis (*S* and *S*′)

If apoptotic cells are not consumed by macrophages, they will, eventually, undergo necrosis. Lipid released from apoptotic cells into the extracellular space contributes to the lipid content of the necrotic core. We assume that apoptotic cells undergo postapoptotic necrosis at rate *ν* (per unit time) such that:

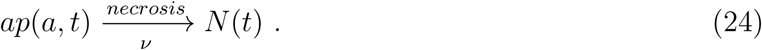

Thus the necrosis sink term *S* in equation (4), is given by:

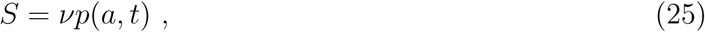

and the source term *S*′ in equation (5) is:

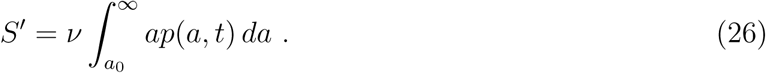

#### 2.2.7 Macrophage ingestion of necrotic material (*C* and *C*′)

We assume that the rate at which a macrophage ingests lipid from the necrotic core is proportional to the amount of material in the core and independent of the macrophage lipid load. Hence, in equation (3),

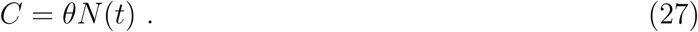

The total rate at which the macrophage population ingests necrotic material is proportional to the total number of macrophages and the amount of necrotic material and, so, in equation (5) we define:

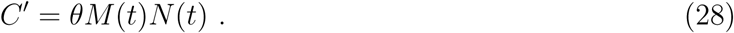

We remark that the functional forms used to define *C* and *C*′ ensure conservation of mass of necrotic material.

### 2.3 Model statement

Having specified functional forms for the cellular processes in our model, we now state the governing equations in full. Substituting from equations (12), (16), (18), (19), (21), (22), (23), (25), (26), (27) and (28) into equations (3)-(5), we obtain the following equations for the time evolution of the number density of live macrophages *m*(*a, t*), apoptotic macrophages *p*(*a, t*) with lipid content *a*_0_ *< a < ∞* and lipid content of the necrotic core *N*(*t*) at time *t* > 0:

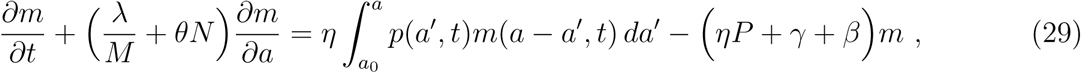

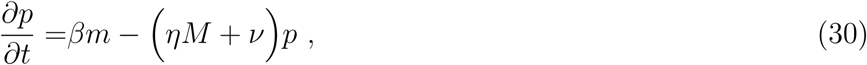

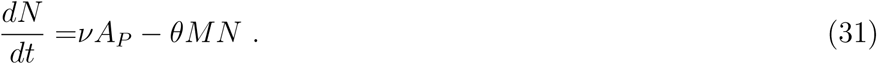

with boundary condition at *a* = *a*_0_:

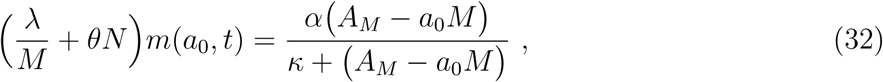

and initial conditions:

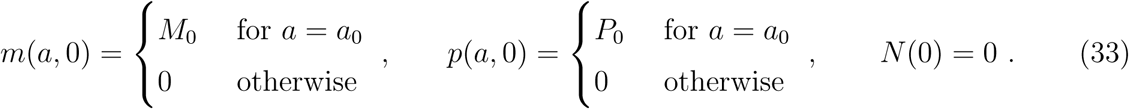

In equations (33), *M*_0_ and *P*_0_ (*M*_0_ *> P*_0_) denote the initial numbers of live and apoptotic macrophages respectively, and we assume that the tissue is initially devoid of necrotic material. By integrating equations (29) and (30) with respect to *a* and assuming that *m*(*a, t*) → 0 and *p*(*a, t*) → 0 as *a → ∞*, we derive the following ordinary differential equations for the time evolution of the total number of live macrophages *M*(*t*) and apoptotic macrophages *P*(*t*) together with their respective initial conditions::

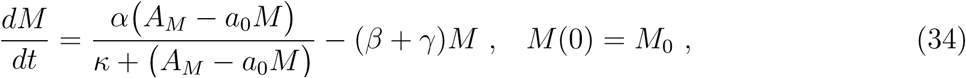

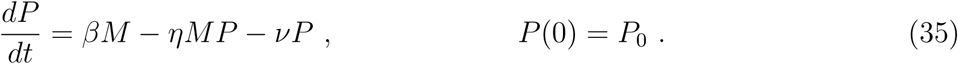

In what follows, it will be useful to keep track of the total quantity of lipid contained by live macrophages *A_M_*(*t*) and apoptotic macrophages *A_P_*(*t*). Ordinary differential equations for *A_M_*(*t*) and *A_P_*(*t*) can be found by multiplying equations (29) and (30) by *a* and integrating with respect to *a*. Making the additional assumption that *am*(*a, t*) → 0 and *ap*(*a, t*) → 0 as *a → ∞* we get:

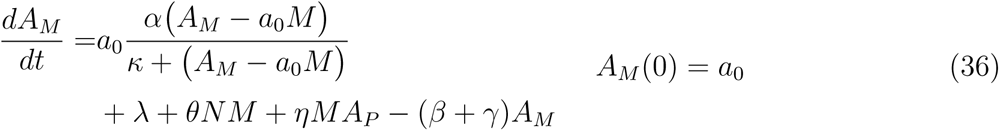

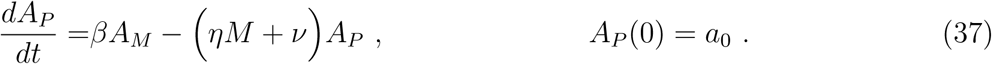

Equations (31) and (34) to (37) together with their associated initial conditions are a closed system of ODEs for *M*, *P*, *A_M_*, *A_P_* and *N*. These give information about the total macrophage and apoptotic cell population size and total lipid loads as well as the size of the necrotic core. The PDEs (29) and (30) for *m*(*a, t*) and *p*(*a, t*) provide detailed information about the distribution of lipid loads across the population of live and apoptotic macrophages. From this distribution it is possible to extract information pertaining to the degree of pathology of the plaque.

### 2.4 Nondimensionalising the model

We recast our model in dimensionless variables. We nondimensionalise time *t* by *β^−^*^1^, the average macrophage lifespan and nondimensionalise lipid content *a* with *a*_0_, the endogenous lipid content of each cell and denote dimensionless quantities with tildes, so that:

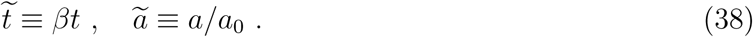

We rescale the dependent variables as follows:

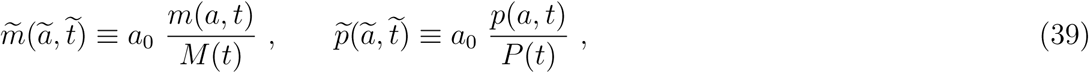

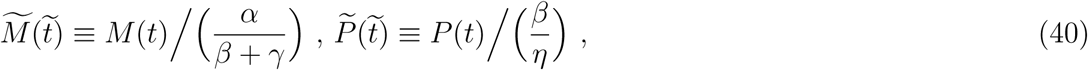

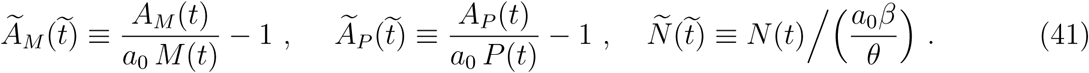

such that 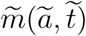 and 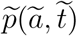 are probability density functions:

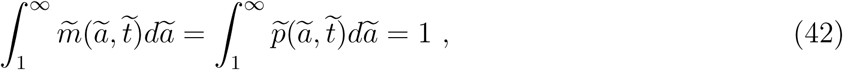

and

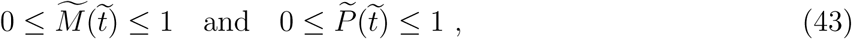

and 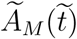 and 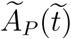 represent the average accumulated lipid content per live and dead macrophage respectively:

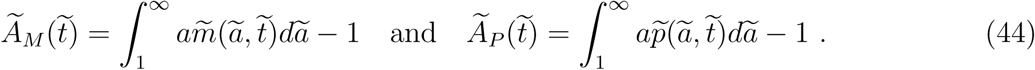

Under these rescalings equations (29) and (30) transform to give the following equations for 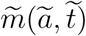 and 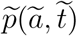 respectively:

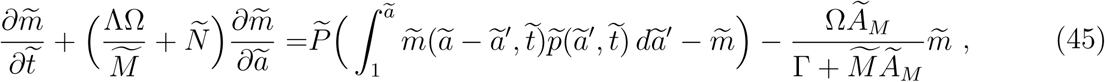

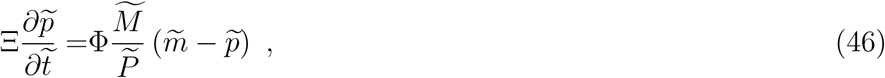

with

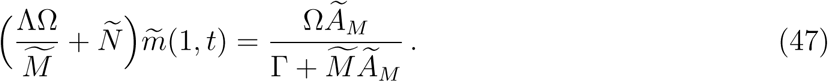

Equations (31), (34), (35), (36) and (37) transform to give the following closed system of ordinary differential equations for 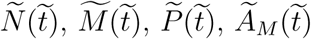 and 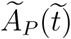 respectively:

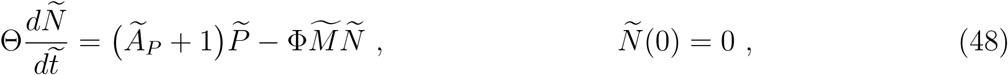

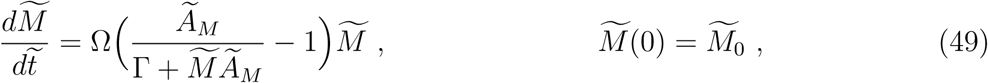

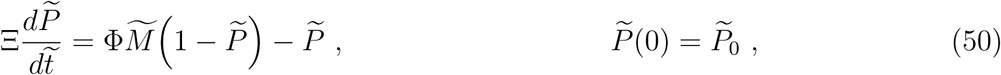

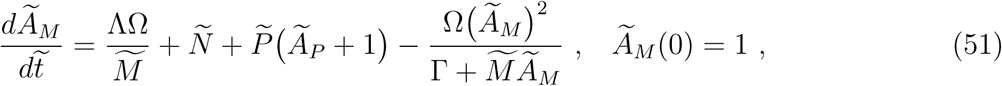

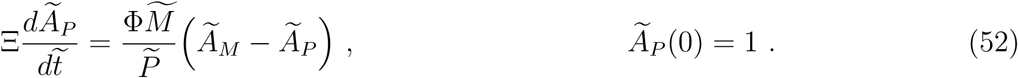

where 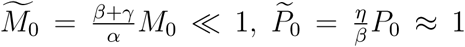 and we have introduced the following dimensionless parameters:

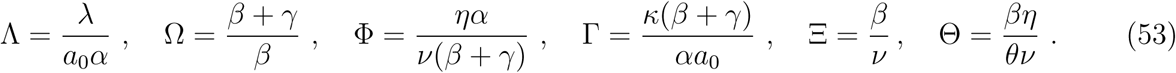

Parameters Λ, Φ, Ω and Γ play key roles in the analysis in the rest of the paper and have the following physical interpretation:

- Parameter Λ *≥* 0 is the ratio of the rate of influx of LDL-derived (exogenous) lipid into the macrophage population to the maximum rate of influx of monocyte-derived (endogenous) lipid into the macrophage population. If 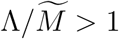 then we anticipate that most of the lipid inside the macrophages derives from LDL; conversely if 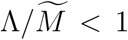 then most internalised lipid derives from apoptotic cells.
- The parameter Ω *≥* 1 governs the proportion of macrophages that undergo apoptosis (at rate *β*) versus emigration out of the plaque (at rate *γ*). The proportion of macrophages that have an apoptotic fate is 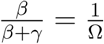 and the proportion which emigrate is 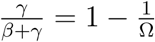.
- The model assumes that emigration and apoptosis are both independent of lipid load. Hence, the average lipid content of the apoptotic cell or emigrating cell is given by the average macrophage lipid content 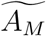 which reflects the rate of LDL, necrotic debris and apoptotic cell ingestion. A macrophage that dies via apoptosis produces one apoptotic cell that is consumed by a live macrophage via efferocytosis (rate *ηM*) or deposits its lipid content into the necrotic core via postapoptotic necrosis (rate *ν*). Thus the proportion of apoptotic macrophages that undergo efferocytosis is 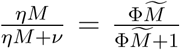 and the proportion that undergo necrosis is 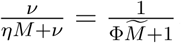.
- The dimensionless parameter Γ governs the rate of saturation of the monocyte recruitment rate function. It represents the quantity of accumulated lipid in the macrophage population when the monocyte recruitment rate is half maximal. For simplicity, without loss of generality we fix Γ = 5 for the remainder of this study.

The dimensionless parameters Ξ and Θ relate to the rate of change of the apoptotic cell population size 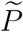 and lipid content 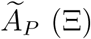 and the rate of change of the necrotic core lipid content 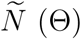. These parameters indicate how quickly the dynamics of 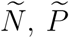 and 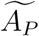 change relative to 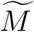 and 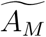. Note that Ξ and Θ are not independent and so changing the rate of evolution of 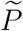 or 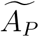, for example, will affect the rate of evolution of 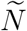. The parameters Ξ and Θ do not affect the long time behaviour of the system.

In the next section, we will omit the tildes and use unadorned symbols to denote nondimensional variables for the remainder of the paper.

### 2.5 A numerical simulation of the time-dependent model

Time-dependent solutions to the model were obtained. Finite difference approximations were used to generate numerical solutions to the governing equations. The midpoint method was used to approximate equations (48)-(52) [41]. An upwinding scheme was used to solve equations (45) and (46), which are first order hyperbolic partial differential equations [41].

To our knowledge, parameter values associated with atherosclerotic plaques *in vivo* are not currently known and may vary markedly between individuals or even plaques. Here, guided by the parameter estimates from [38], biologically plausible parameters values were chosen to simulate four different plaque states illustrated in Figure 2. The chosen values and reasoning for each state are as follows. To model a low-grade inflammatory state we chose a parameter set for which the lipid gain per cell via LDL is slightly higher than the lipid removal per cell via HDL (Λ = 0.01), macrophages are ten-times more likely to undergo apoptotic death than emigration from the plaque (Ω − 1 = 0.1) [8] and the rate of efferocytosis is high relative to the rate of necrosis so that Φ = 100 which implies that postapoptotic necrosis is negligible. To model a severe inflammatory response to elevated LDL levels in the serum, we increased Λ from Λ = 0.01 to Λ = 10 at time *t* = 200. To model the onset of plaque pathology associated with reduced efferocytosis we reduced Φ from Φ = 100 to Φ = 0.1 at time *t* = 400. To model plaque regression, we increased Ω from Ω = 1.1 to Ω = 2 to reflect either an increased emigration rate and/or decreased apoptosis rate. Note that only one parameter is change in value at a time; Φ and Ω hold their value when Λ increases in value at *t* = 200, etc.

**Figure 2:**
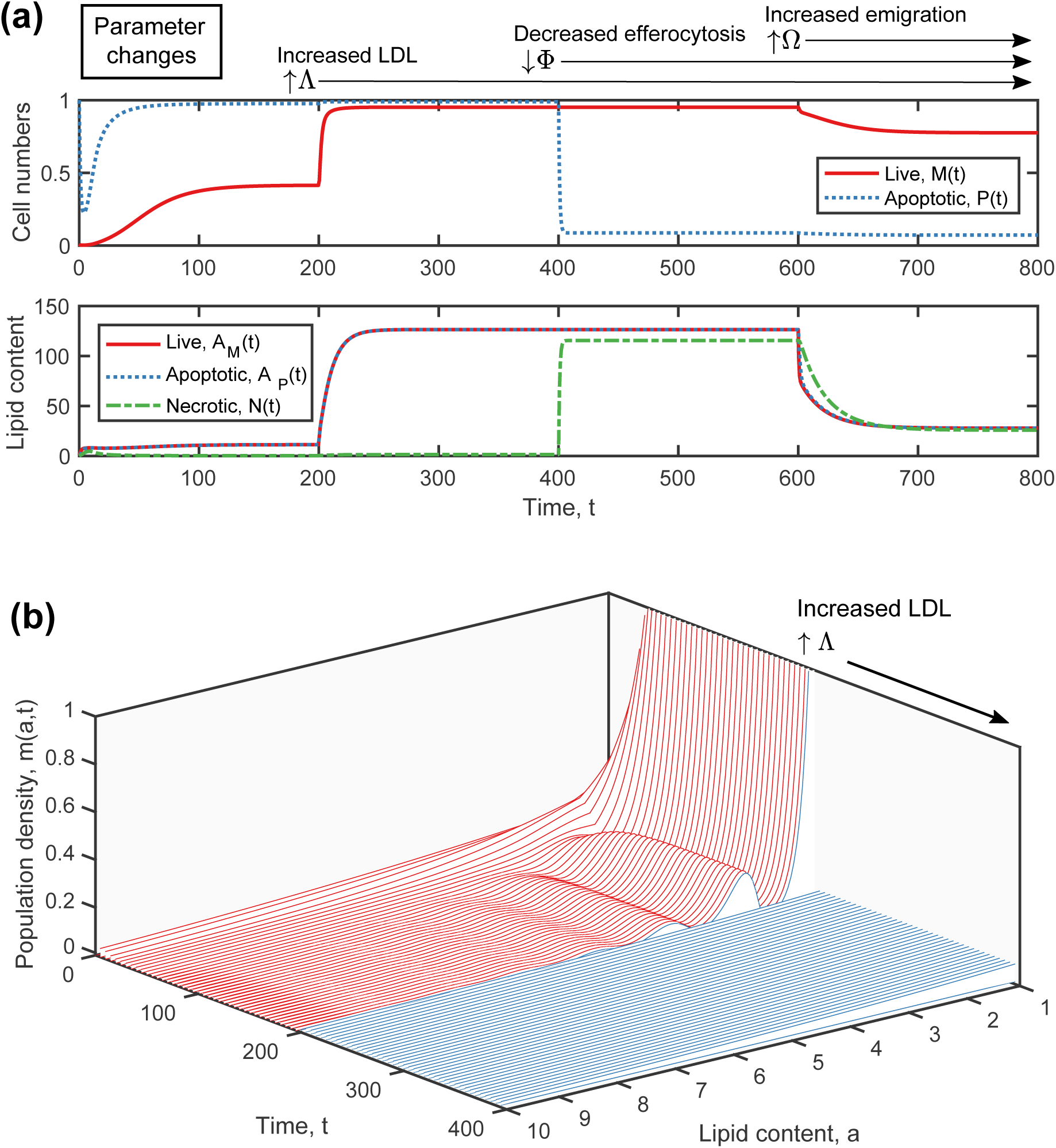
Numerical Results. (a) Time evolution of the number cells in the population, both live *M*(*t*) and apoptotic *P*(*t*) (top) and average accumulated lipid content of live macrophages *A_M_*(*t*) and apoptotic macrophages *A_P_*(*t*) and the lipid content of the necrotic core *N*(*t*) (bottom). We increase the influx of LDL from Λ = 0.01 to 10 at *t* = 200; we decrease the rate of efferocytosis from Φ = 100 to 0.1 at *t* = 400; and increased the rate of cell emigration from Ω = 1.1 to Ω = 2 at *t* = 600. Only one parameter is changed at a time and all other parameters are held fixed at each change and Γ = 5, Ξ = *θ* = 1 throughout. (b) The number density of live macrophages *m*(*a, t*) as a function of both accumulated lipid *a* and time *t*. This result is derived from 0 *≤ t ≤* 400 in the simulation shown in (a).

Figure 2 illustrates results from a numerical simulation of the nondimensional model given by equations (45)-(52). As outlined above, parameters Λ, Φ and Ω change at times *t* = 200, 400 and 600 respectively while parameters Γ, Θ and Ξ are held constant. The top two plots of Figure 2 show how the number and average accumulated lipid content of live and apoptotic macrophages change over time and also how the lipid content of the necrotic core evolves. High rates of LDL ingestion rate, modelled by high values of Λ, lead to the appearance of large numbers of foam cells which drive rapid monocyte recruitment into the tissue. Low efferocytosis rates (that is, low values of Φ) promote postapoptotic necrosis and produce a necrotic core. Together, high rates of LDL ingestion and low efferocytosis rates produce large numbers of postapoptotic foam cells and a large lipid-rich necrotic core. High macrophage emigration rates, (high values of Ω) remove large quantities of lipid from the system as lipid-loaded cells leave the plaque and, consequently, reduce lipid accumulation inside macrophages and the necrotic core.

The bottom plot in Figure 2 shows changes in the lipid distribution across the macrophage population (*m*(*a, t*)). For high rates of LDL ingestion, lipid is monotonically distributed across the macrophage population. In this case, the extent of lipid accumulation and population heterogeneity with respect to lipid content is large. By contrast, when LDL ingestion rates are low, the lipid distribution is principally determined by phagocytosis of apoptotic cells and, as a result, has a series of peaks, separated approximately by the amount of endogenous lipid in each cell.

In Figure 2 the timescale of interest is the average lifespan of a macrophage before it undergoes apoptosis. We note, for the parameter values used here, that the system approaches a steady state solution generally within a non-dimensional time span of 100 macrophage lifespans or less. In inflamed tissues, the macrophage lifespan is on the order of days to weeks [6, 8] and so 100 macrophage lifespans might correspond to 0.5-2 years. Since plaques can exist for decades, this suggests that it may be sufficient to study the steady state distribution of lipid in the live and apoptotic macrophage populations.

## 3 Analysis and results

We assume that lipid loads, the numbers of live and apoptotic macrophages, and the lipid in the necrotic core reach a steady state quickly, relative to the lifetime of the plaque and in the absence of physiological changes. These changes might include an increased rate of LDL influx into the plaque or decreased rate of efferocytosis. We examine the steady state solutions of equations (45) to (52) where all time derivatives are set to zero. We first focus on the ODE system defined by equations (48) to (52).

### 3.1 Steady state cell populations and lipid content

#### 3.1.1 Steady states of the ODE system

We assume that as *t → ∞* the variables *M*(*t*), *P*(*t*), *N*(*t*), *A_M_*(*t*) and *A_P_*(*t*) approach steady state values *M\**, *P\**, *N\**, 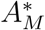 and 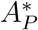 respectively. Setting time derivatives to zero in equations (48)-(52), we deduce that *M\**, *P\**, *N\** and 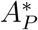 can be defined in terms of 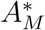 as follows:

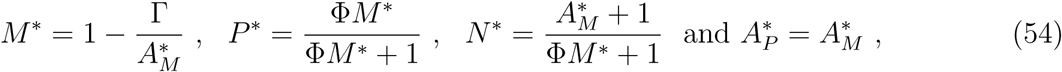

where 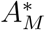 satisfies the following quadratic equation:

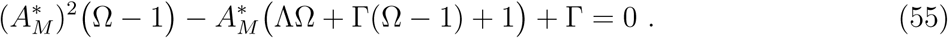

Equations (54) and (55) are used to plot *M\**, *P\**, 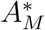, 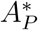 and *N\** in Figure 3 for various values of parameters Λ, Ω and Φ and fixed Γ = 5.

**Figure 3:**
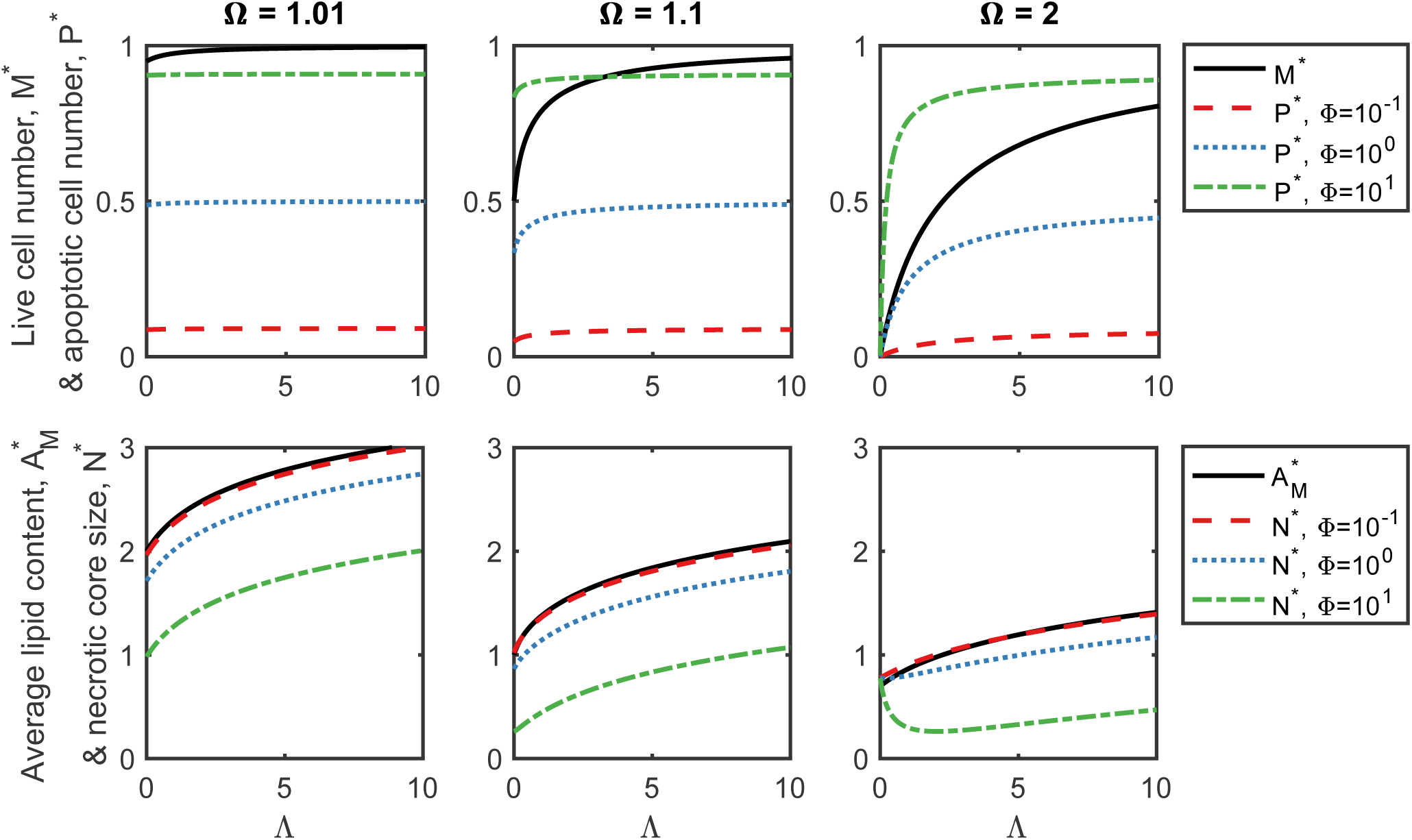
**Long time population sizes and lipid contents** calculated from equations (54) and 55). The live cell numbers *M\** (black line; top row), average accumulated lipid content 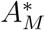 (black line; bottom row), apoptotic cell numbers *P\** (coloured dots and dashes; top row) and lipid content of the necrotic core *N\** *(coloured dots and dashes; bottom row) for a range of Λ (x-axis), Ω (left to right plots) and Φ (colours). Here we have fixed Γ = 5.

For Λ > 0 and Ω > 1, equation (55) admits two real positive solutions: one solution has 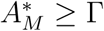 and the other 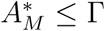. Since we require *M* ≥* 0 for physically real solutions, we reject the solution with 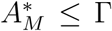. In the special case when Λ = 0, there are two solutions: 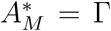 when 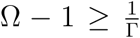 or 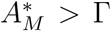 when 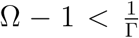. The first solution gives *M\** = 0 which implies that the immune response does not exist when there is no net LDL ingestion and large numbers of macrophages emigrate from the tissue. As Ω → 1 (no emigration) then the real solution 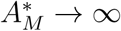 and *M* → 1 because without net lipid efflux from cells or cell emigration there is no mechanism for lipid to leave the plaque and as a result, it progressively accumulates inside the macrophage population.

Physically unrealistic (i.e. complex and/or negative) solutions may exist when Λ < 0; that is, when HDL exports lipid from the cells at a faster rate than that at which it accumulates due to LDL ingestion. In this case, cells are likely to have low lipid loads and so macrophage recruitment and macrophage numbers will remain at low levels. This case, which represents healthy tissue, is not of primary interest in this study and so we will henceforth assume that Λ > 0.

#### 3.1.2 Pathological states

In our model, inflammation resolves only when lipid gained via LDL ingestion is less than lipid removed via HDL (Λ ≤ 0) and when at least 20% (if Γ = 5) of all macrophages emigrate from the tissue (as opposed to die in the tissue). Otherwise, a nonresolving inflammatory response occurs.

An atherosclerotic plaque is considered pathological and potentially harmful when it contains large numbers of macrophages (say *M\** > 0.9) with large lipid loads (say 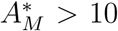) and a large necrotic core (say *N\** > 10) [13, 42, 43]. Suppose that the plaque is non-pathological/benign when *M\**, 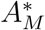 and *N\** are all small. The pathological state arises in our model for large LDL influx rates into the tissue (say Λ > 1), low emigration rates (say Ω − 1 < 0.2) and ineffective efferocytosis (say Φ < 10). In this state, a large necrotic core forms when large numbers of macrophage foam cells die via apoptosis and then undergo postapoptotic necrosis as a result of defective efferocytosis. The non-pathological state in our model occurs only for low LDL influx rates (Λ < 1), high emigration rates (Ω − 1 > 0.2) and high efferocytosis rates relative to the rate of secondary necrosis (Φ > 10). In this state, macrophages clear LDL, apoptotic cells and necrotic debris and ferry the lipid out of the plaque via cell emigration and HDL export. There are several inflammatory states which are not necessarily pathological. For example, for large LDL influx rates, low emigration rates and high efferocytosis rates, there can be large numbers of foam cells (large *M\** and 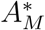) without the presence of a necrotic core (small *N\**) as efficient efferocytosis blocks postapoptotic necrosis. By contrast, for low efferocytosis rates, high LDL influx rates and high emigration rates the necrotic core remains small because high macrophage emigration rates prevents apoptosis and hence postapoptotic necrosis.

### 3.2 Steady state lipid distribution in the macrophages

### 3.2.1 Governing equation

We assume that as *t → ∞* the variables *m*(*a, t*) and *p*(*a, t*) approach steady state distributions *m\**(*a*) and *p\**(*a*). From equations (45) and (46), with time derivatives set to zero 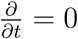, we deduce that *p\**(*a*) = *m\**(*a*) where *m\**(*a*) solves the following ordinary integro-differential equation:

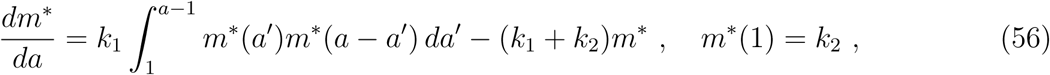

where

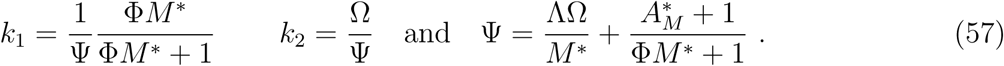

Note that *k*_1_ and *k*_2_ are functions of *M\** and 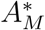 which also depend on model parameters Φ, Ω and Λ (equations (54) and (55)). Parameter Ψ is the mean rate of influx of lipid into each macrophage via LDL and via phagocytosis of necrotic debris. We note that *k*_1_ *≤ k*_2_ as 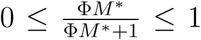 and Ω *≥* 1.

**Table 1:** Summary of qualitative behavious exhibited by our mathematical model as the LDL influx rate, macrophage emigration rate and efferocytosis rate vary.

| $\Lambda$ | $\Phi$ | $\Omega$ | |
| --- | --- | --- | --- |
| LDL influx rate | rate of efferocytosis relative to necrosis | rate of emigration relative to apoptosis | Model prediction |
| High | Low | Low | Large necrotic core, large number of macrophages in the plaque, a highly pathological state |
| High | Low | High | A large number of macrophages, but small necrotic core. Lipid is ferried out of the plaque by high rate of macrophage emigration. |
| High | High | Low | A large number of macrophages heavily laden with lipid, but small necrotic core. Not intrinsically dangerous but vulnerable to changes in parameter values which would drive the plaque to pathology. |
| Low | High | High | Small necrotic core, low numbers of macrophages, nonpathological state. |

Contour plots showing how *k*_1_ and *k*_2_ change as the parameters Λ, Ω and Φ vary are shown in Figure 4. The functions *k*_1_ and *k*_2_ depend on Λ, Ω and Φ in broadly similar ways. Both *k*_1_ and *k*_2_ take on their highest values when Λ and Ω are low and Φ is high. For fixed values of Φ, both *k*_1_ and *k*_2_ have maxima in Ω between Ω = 1 and Ω = 1.2 for Λ = 0.1 and 1.

**Figure 4:**
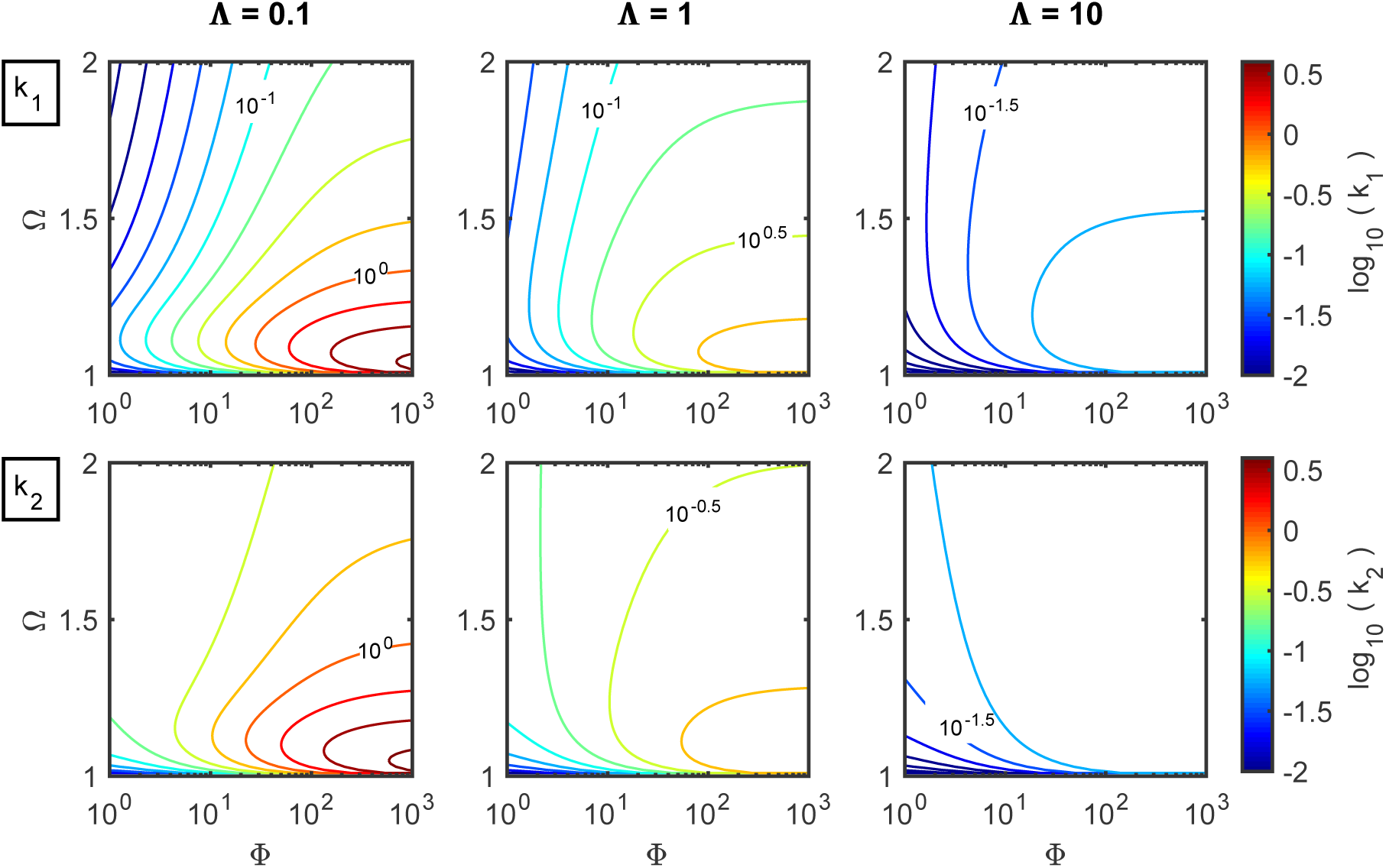
Contour plots showing how the parameter groupings *k*_1_ (top row) and *k*_2_ (bottom row) as defined by equation (57) change as the model parameters Ω (y-axis), Φ (x-axis) and Λ (columns) are varied. As before, we set Γ = 5. The parameter groupings *k*_1_ and *k*_2_ govern how lipid is distributed in the steady state macrophage population, *m\**(*a*).

#### 3.2.2 Analytic solution for steady state macrophage population as a function of lipid load

An analytical solution for *m\**(*a*) can be constructed by solving equation (56) recursively on successive unit intervals *n ≤ a < n* + 1, *n ∈* ℕ, *n ≥* 1. Let *m_n_*(*a*) be the solution on the interval *n ≤ a < n* + 1. Then, equation (56) can be recast as a system of integro-differential equations, each defined on a particular interval with *m_n_*(*n* + 1) = *m_n_*_+1_(*n* + 1) to ensure continuity of the solution. For 1 *≤ a <* 2, 2 *≤ a <* 3 and 3 *≤ a <* 4, we have respectively:

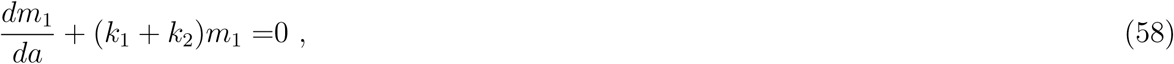

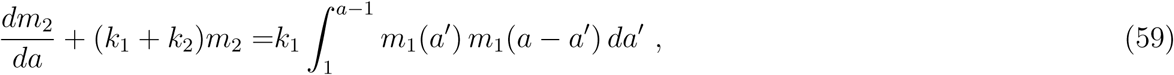

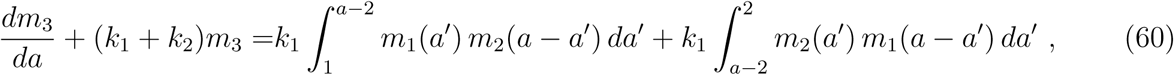

where *m*_1_(1) = *k*_2_, *m*_2_(2) = *m*_1_(2) and *m*_3_(3) = *m*_2_(3). In general, for *n ≤ a < n* + 1, we have:

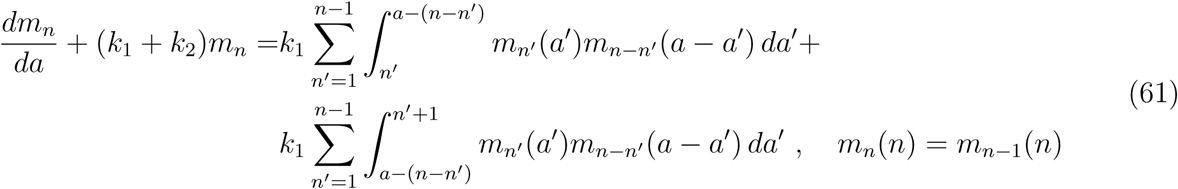

We note that there is no integral term in equation (58) since all live apoptotic macrophages have lipid load *a ≥* 1. Hence the minimum combined lipid load of a live macrophage and the apoptotic cell that it ingests is two. Integration of equation (58) supplies:

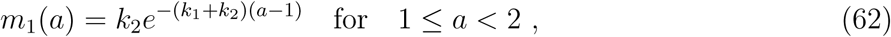

whilst equations (59) and (60) yield:

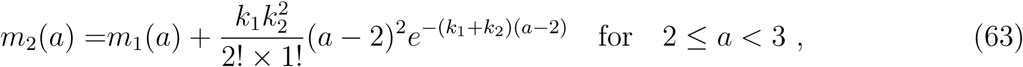

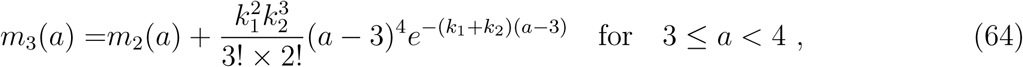

and more generally:

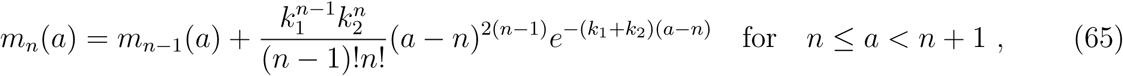

We deduce finally that solution *m\**(*a*), the normalised steady state lipid distribution, can be written as:

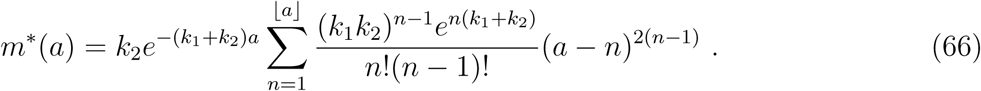

This solution is plotted in Figure 5 for the parameter values used to generate the numerical simulation shown in Figure 2.

**Figure 5:**
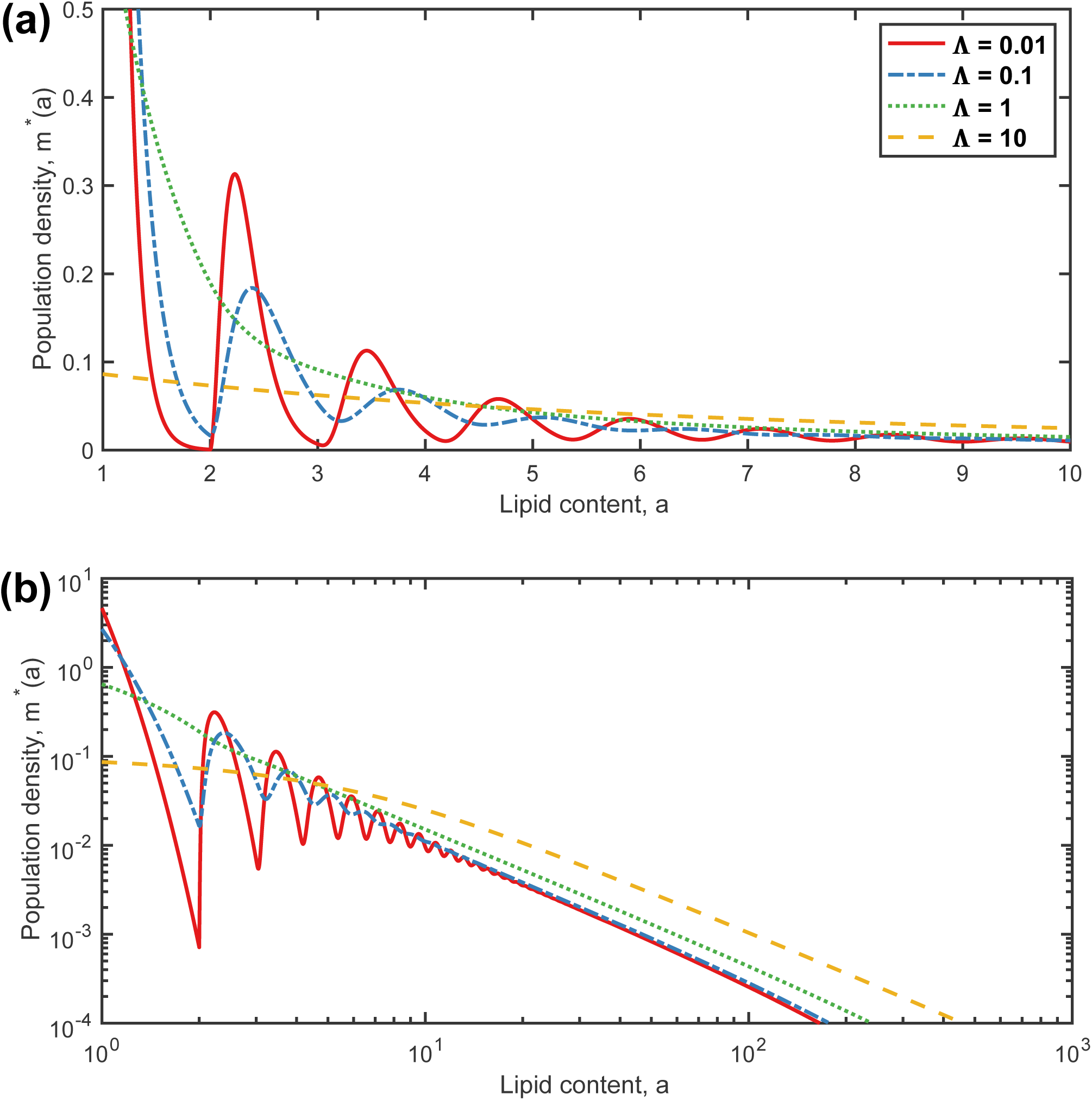
Analytical results showing how the proportion of both live and apoptotic macrophages (*m\**(*a*) = *p\**(*a*) as defined by equation (65)) changes with lipid content *a*, at steady state, for different values of the LDL influx: Λ = 0.01 (red solid line), 0.1 (blue dash-dotted line), 1 (green dotted line) and 10 (yellow dashed line). The emigration rate relative to apoptosis rate is fixed at Ω = 1.1 and the efferocytosis rate relative to postapoptotic necrosis rate at Φ = 100. These parameter values were used in the numerical results shown in Figure 2. Both plots show how the steady state solutions vary with *a*; the upper plot is on linear axes while the lower plot is on a log-log scale.

There are two key qualitative properties of *m\**(*a*) evident in Figure 5. First, as Λ increases, the profile of *m\**(*a*), particularly for small *a*, changes from having multiple local maxima to being monotonic. Second, for large *a* the profile decays algebraically as *a → ∞* at the same rate, regardless of the value of Λ.

#### 3.2.3 Two qualitatively different profiles

In Figure 5 the profile of *m\**(*a*) either monotonically decreases or attains a series of local maxima. Analysis of the local maximum and minimum on 2 *≤ a <* 3 enables us to identify the parameter regimes in which *m\**(*a*) is either peaked or monotonic. We define the lipid content of the maximum and minimum values on 2 *≤ a <* 3 as *a ≡ a*_+_ and *a ≡ a_−_* respectively such that 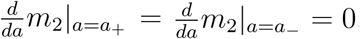. From equation (63) we find that:

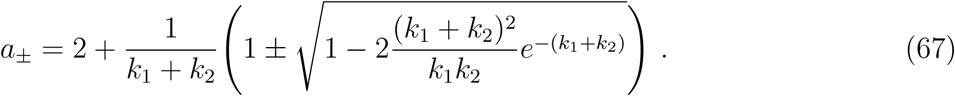

The local maximum and minimum in the region 2 *≤ a <* 3 coalesce when *a*_+_ = *a_−_* and this occurs when:

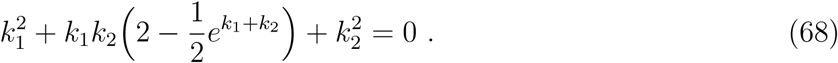

In Figure 6 we show how *a_±_* change as *k*_1_ and *k*_2_ vary, remembering the dependence of *k*_1_ and *k*_2_ on Λ, Ω and Φ depicted in Figure 4.

**Figure 6:**
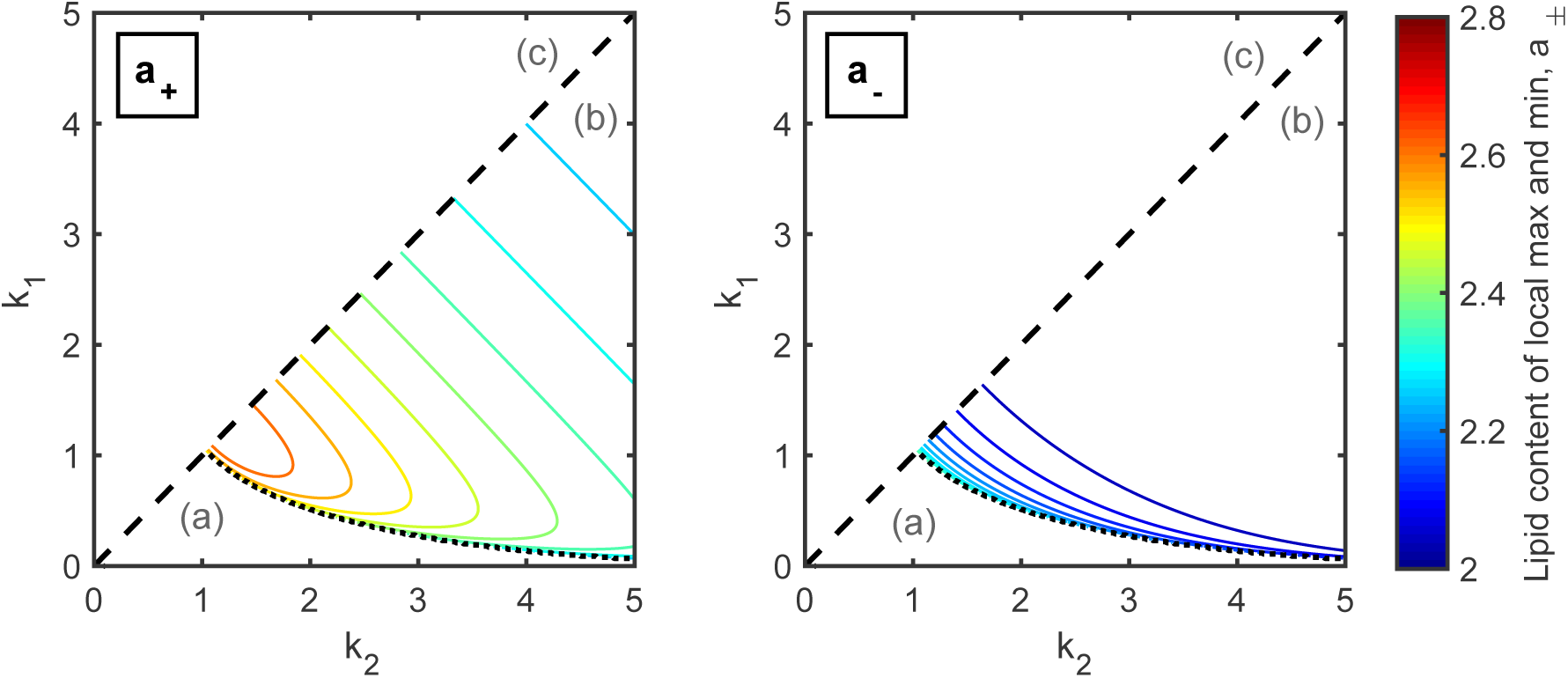
Analytical results showing the lipid content of the local maximum (*a*_+_) and minimum (*a_−_*) value of the steady state solution *m\**(*a*) across 2 *≤ a <* 3 for various *k*_1_ and *k*_2_ values (equation (57)) on a contour plot. Here *a*_+_ and *a_−_* are determined by equation (67). The dotted line is the solution where *a*_+_ = *a_−_* given by equation (68) which separates parameter space into regimes where either *m\**(*a*) is monotonic (region (a)) or peaked (region (b)) profile. This dotted line can be crudely approximated as *k*_1_ *≈* 1*/k*_2_. The dashed line where *k*_1_ = *k*_2_ separates infeasible regions (*k*_1_ *> k*_2_; labelled (c)) from feasible ones (*k*_1_ *≤ k*_2_). An explanation why *k*_1_ *≤ k*_2_ is given beneath equation (57).

Figure 6 shows that *m\**(*a*) has a peaked profile for *k*_2_ > 1, *k*_2_ *> k* and *k*_1_ ⪸ 1*/k*_2_ (region (b) in Figure 6). Figure 4 shows that *k*_2_ > 1 when the LDL influx rate into the tissue is small (Λ < 1), the cell emigration rate relative to the apoptosis rate is small (Ω < 1.1) and the efferocytosis rate relative to necrosis rate is large (Φ > 10). That is during typical inflammatory responses where net lipid uptake from LDL ingestion is negligible, when most macrophages die via apoptosis and most apoptotic cells are consumed, the macrophage lipid load may have a single peak in each interval *n, a < n* + 1.

Figure 6 shows that *m\**(*a*) has a monotonic profile when 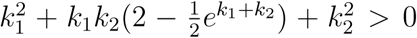(region (a) in Figure 6). These parameter values may correspond to either pathological or healthy states. Figure 4 shows that *k*_1_ < 1 for large Ω which represents a resolving inflammatory response where most macrophages emigrate from the tissue and macrophage recruitment is low so that there are small numbers of macrophages (*M\**) with a small average lipid content (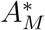). Figure 4 shows that *k*_1_ < 1 also for pathological inflammatory responses where LDL ingestion (Λ) is large so that there are large numbers of macrophages (*M\**) with a large average lipid content (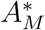). In this case, Ω can take on a wider range of values, but Φ remains low. The low value of Φ could be attributed to low rates of efferocytosis relative to necrosis which leads to the formation of a large necrotic core. In summary, our model shows that the steady state distribution of lipid across macrophages is closely linked to plaque pathology.

### 3.3 Special case: no LDL ingestion or lipid-offloading to HDL (Λ = 0)

Macrophage foam cell formation via extensive LDL ingestion is a unique feature of atherosclerotic plaques. However, macrophage foam cells also commonly appear in chronic lesions that do not contain modified LDL [10] Here, we show that efferocytosis alone, with no net ingestion of LDL or lipid export to HDL, can cause macrophages to accumulate lipid derived from the membranes of macrophages that have died and been consumed.

We suppose that there is no net LDL influx into the tissue and no net lipid export from macrophages, so that Λ = 0. We further assume that the rate of phagocytosis (given by *ηM* in the dimensional equations) is more rapid than the rate of apoptosis (*β*) and secondary necrosis (*ν*) such that *η » ν, β* which, in terms of the nondimensional variables, implies Φ*M »* 1. We assume further that every apoptotic macrophage is consumed so that in the limit as Φ*M* → ∞ (given *M* > 0). If Φ*M* → ∞ then the nondimensional equations (46), (50) and (52) become:

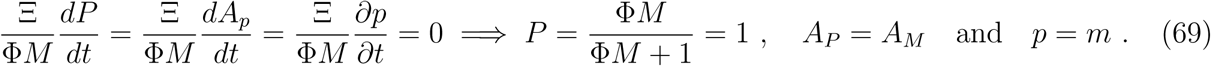

Under this assumption apoptotic cells are ingested rapidly by live macrophages soon after they have undergone apoptosis. With these relations, equations (45), (49) and (51) become:

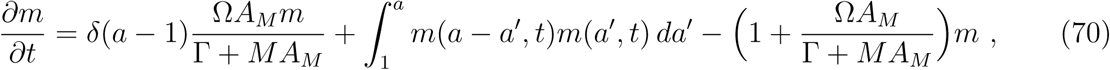

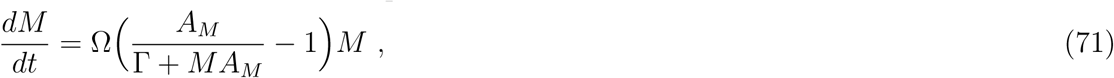

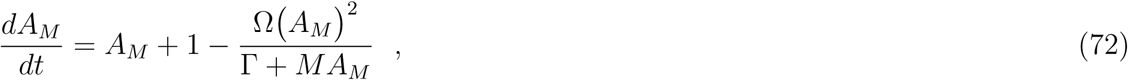

where *δ*(*a −* 1) is the delta function. Here we have incorporated the boundary condition (47) into equation (70) in the same way that the recruitment term was part of the differential equation for *m* in the model setup in Section 2.1 and Subsection 2.2.2.

If macrophages do not ingest LDL, then they will only accumulate lipid in integer units of the endogeneous quantity of lipid, which is *a*_0_ in dimensional terms and 1 in the nondimensional system. (We are assuming here, of course, that all macrophages have an equal quantity of endogeneous lipid.) Hence, on *n* ≤ a < *n* + 1, we set

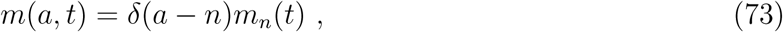

where *m_n_*(*t*) is the proportion of macrophages at time *t* that contain *n* units of the endogenous lipid content. We substitute equation (73) into equation (70) to produce the following Smoluchowski coagulation equation [44]:

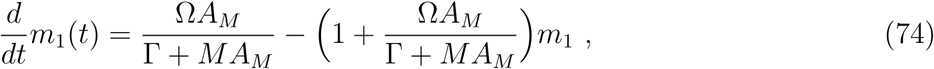

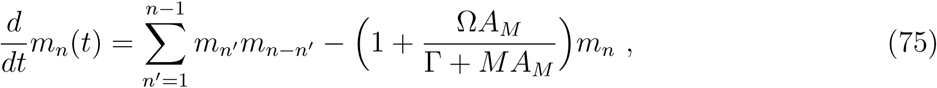

where *A_M_* and *M* are given by equations (71) and (72). The fact that equation (74) and (75) belongs to the family of Smoluchowski coagulation equations implies that efferocytosis causes endogenous lipid to accumulate inside macrophage populations via a coalescence (or coagulation) process [45, 46]

We are primarily interested in the steady state solutions of equations (74) and (75) where 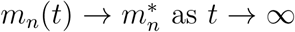. By setting the time derivatives equal to zero in equations (71), (74) and (75), we find that 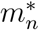 is given by the following recurrence relation:

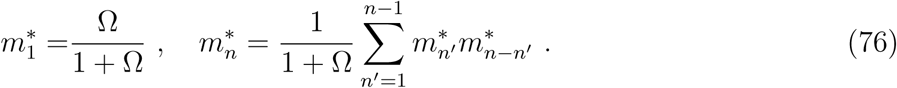

Note equation (76) can also be derived from equation (61) in the limit *k*_1_*, k*_2_ *→ ∞* and *k*_2_*/k*_1_ → Ω and when all derivatives with respect to *a* are equal to zero. By comparing equation (76) to the recurrence relation that generates the Catalan numbers, we deduce that 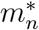 has following solution:

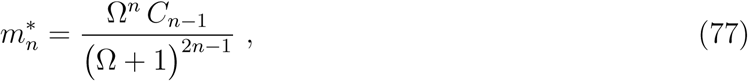

where *C_n_* is the *n*-th Catalan number. In Figure 7 we use equation (77) to plot 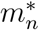 for various values of Ω. In Figure 8 we use equations (66) and (8) to compare the continuous distribution *m\**(*a*) from (66) with the discrete distributions 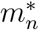 where Λ = 0 from (8). From this figure it is clear that efferocytosis drives, LDL phagocytosis exacerbates and cell emigration reduces excessive lipid accumulation inside macrophages.

Figure 7 shows that for *n »* 1, 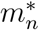 decays algebraically for Ω = 1 and exponentially for Ω > 1. That is, the behaviour of 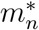 is qualitatively different when Ω = 1 from its behaviour for Ω > 1.

**Figure 7:**
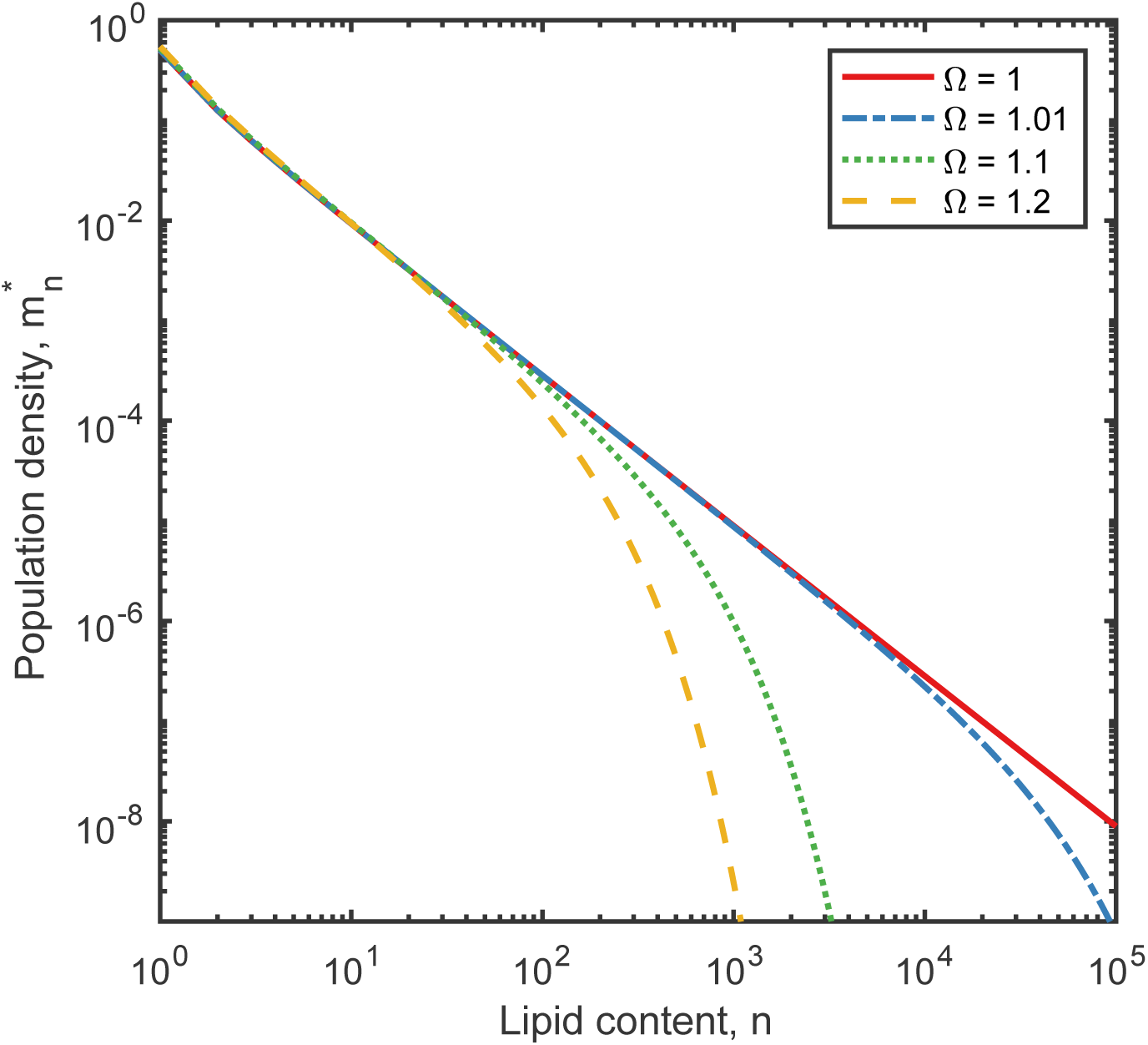
Series of plots showing the proportion of cells 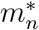 with lipid content *n*, at steady state, when LDL ingestion and lipid offloading are neglected (Λ = 0) for different choices of the emigration rate relative to the apoptosis rate Ω = 1 (red solid line), 1.01 (blue dot-dashed line), 1.1 (green dotted line) and 1.2 (yellow dashed line). This solution was determined analytically from equation (77).

**Figure 8:**
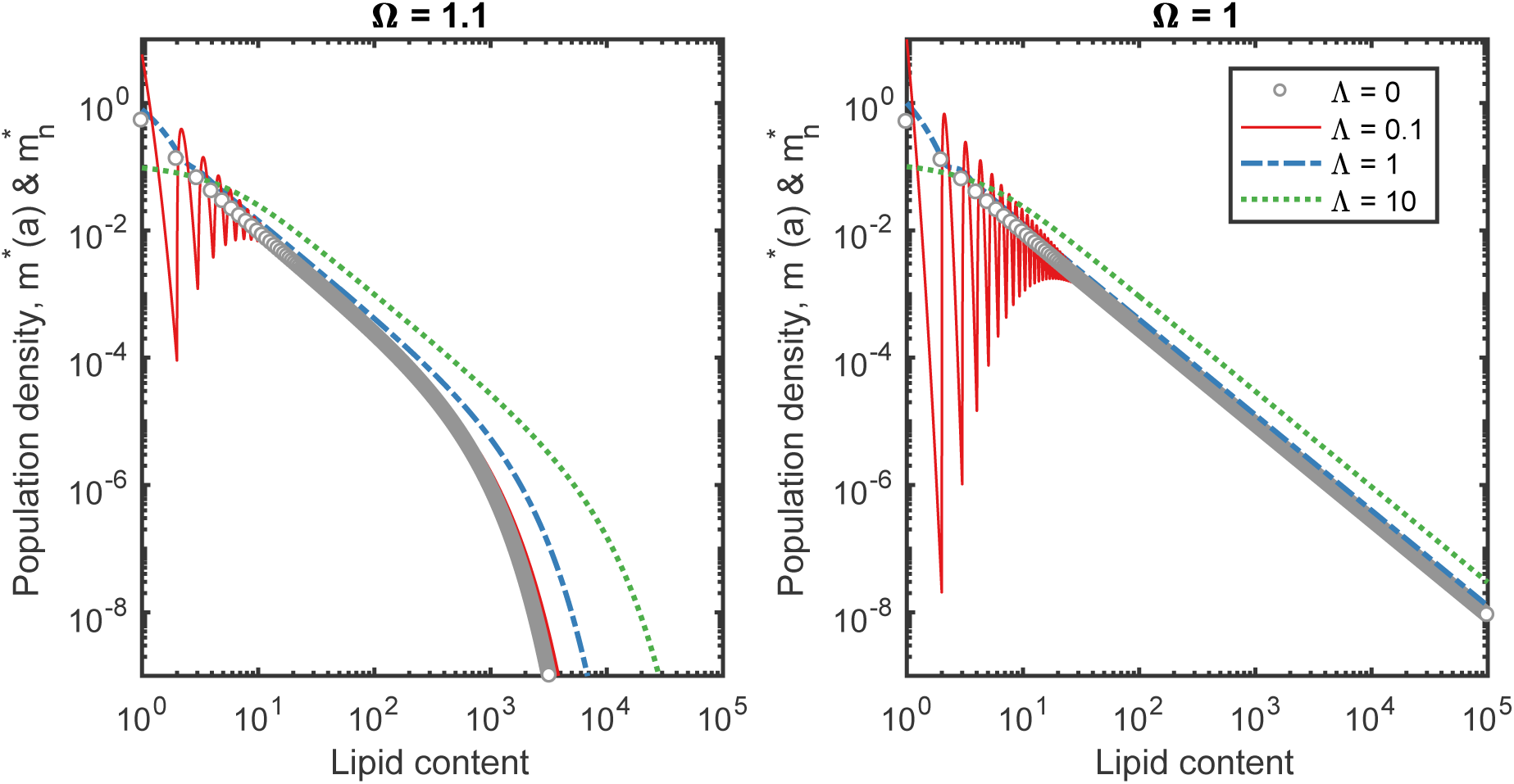
Series of plots showing the steady state proportion of cells 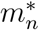 with lipid content *n* without LDL ingestion (Λ = 0, grey circles) and the proportion of cells *m\**(*a*) with lipid content *a* with LDL ingestion (Λ = 0.1 (red solid line), 1 (blue dot-dashed line) and 10 (green dotted line)) with the emigration rate relative to the apoptosis rate Ω = 1.1 (left) and Ω = 1 (right). These solutions were determined analytically from equations (77) and (66).

To understand why the behaviour of 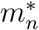 changes at Ω = 1 we recall that *C_n_* = (2*n*!)*/*(*n*!(*n* + 1)!) and use Stirling’s approximation (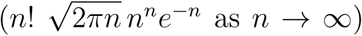) to simplify equation (77). In this way we deduce

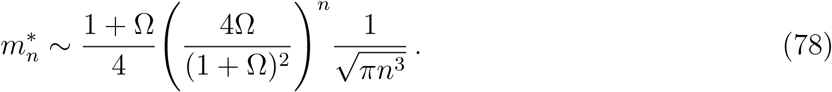

From equation (78) we conclude that, when Ω > 1, the decay in 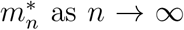 as *n → ∞* is dominated by the exponential term. When Ω = 1 (no cell emigration), 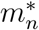 decays algebraically. The qualitative change in *m\**(*a*) as Ω → 1 indicates that only small rates of emigration are required to significantly reduce the coalescence of accumulated lipid, suppress excessive lipid accumulation and remove accumulated lipid from the population.

## Discussion

We have developed and analysed a mathematical model, formulated as a system of integrodifferential equations, that describes how the number and lipid content of live and apoptotic macrophages and the amount of necrotic material in an atherosclerotic plaque change with time. The model couples lipid and inflammation dynamics to show, for the first time, how inflammation shapes lipid accumulation inside macrophages and how this internalised lipid accumulation then shapes inflammation. The model describes how lipid enters, leaves, accumulates and redistributes inside inflammatory monocyte-derived macrophages as these cell enter the tissue, take up LDL and necrotic debris, offload lipid to HDL, die via apoptosis, consume apoptotic cells via efferocytosis and leave the tissue.

By constructing steady state solutions to the governing equations, we are able to assess the impact on plaque formation of known risk factors such as high levels of LDL cholesterol in blood serum, high blood monocyte counts and inefficient phagocytosis by macrophages.

The model developed in this paper is focused on lipid and macrophage dynamics during inflammation and in particular lipid accumulation by macrophages as a consequence of phagocytosis. It considers cellular dynamics within the artery wall rather than the interaction of the artery wall with the bloodsteam or the mechanisms of cell and lipid perfusion through the blood vessel wall. Because this is a simple, analytically tractable, foundational model, we do not include spatial structure as, for example in [29]. Clearly there is scope to include internalised lipid structure in macrophage in many different types models for atherosclerotic plaque development. The model presented here is intended to generate understanding of the role and consequences of this type of macrophage population structure. Ultimately many different models may be combined into more comprehensive models of atherosclerotic plaque development. Models for specific processes and species in the plaque also have the capacity to generate significant new insights on their own and hence are valuable in their own right.

The key insights into atherosclerosis provided by our model are summarised in Figure 9. During typical inflammatory responses, millions of monocyte-derived macrophages turn over daily, primarily via monocyte recruitment (as opposed to cell division), apoptosis (as opposed to cell emigration) and efferocytosis (as opposed to postapoptotic necrosis) [6, 8]. Lipid is introduced into the tissue, not only via LDL influx, but, significantly, via monocyte recruitment which brings cells’ endogenous lipid into the plaque. Lipid that has ben internalised via phagocytosis is transferred from one cell to another via efferocytosis of apoptotic cells.

**Figure 9:**
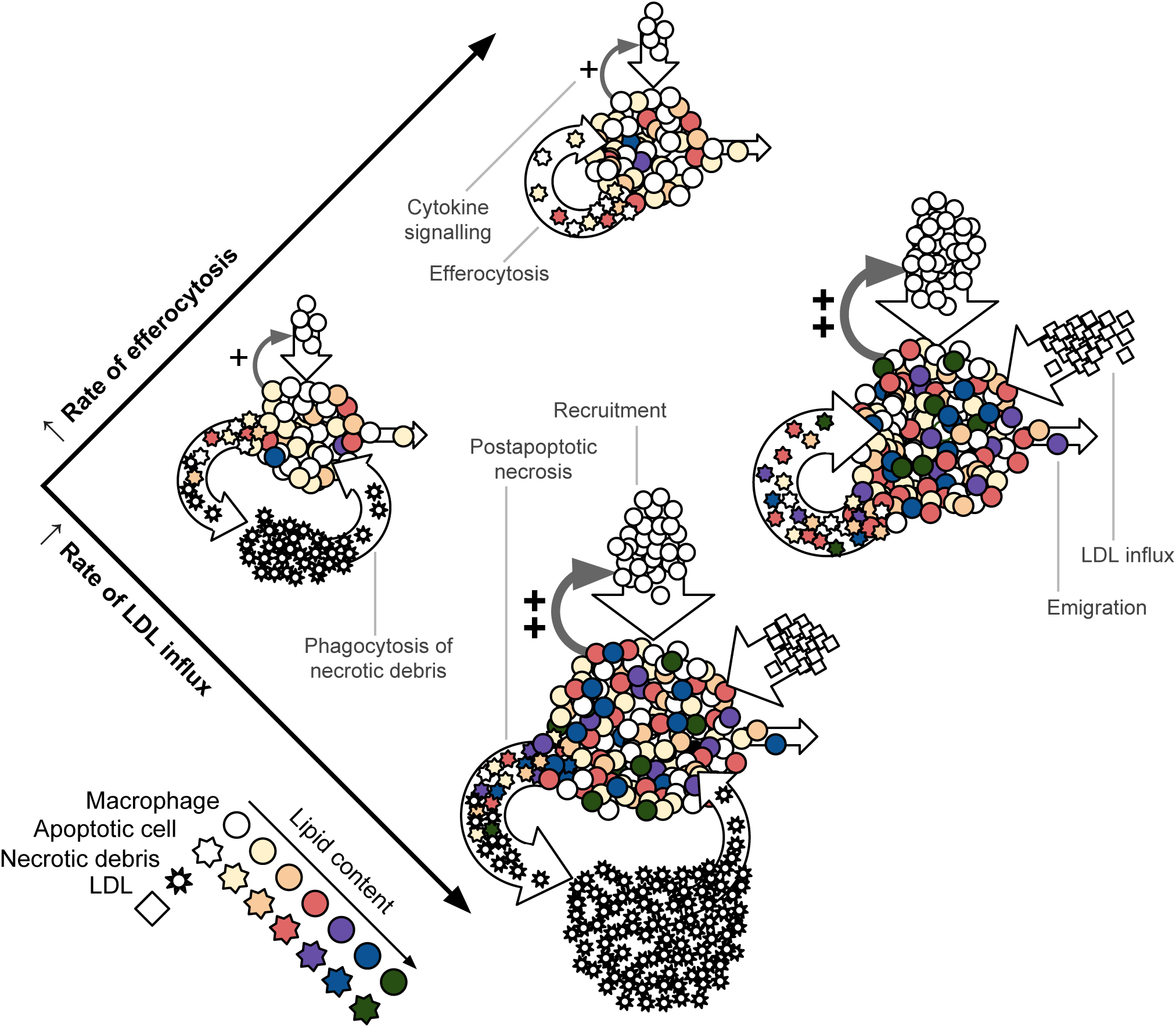
A schematic diagram that summarises the key results derived from our model. Each of the four constituent diagrams represent a different type of plaque. Macrophages are represented by circles, apoptotic macrophages by clouds and necrotic debris by stars. Each plaque experiences macrophage recruitment, which may be amplified by cytokine production by macrophages that are already in the plaque; macrophage emgration; and macrophage apoptosis which is followed either by efferocytosis of the apoptotic cells (top and middle right) or secondary necrosis of the apoptotic cells followed by phagocytosis of the necrotic material (middle left and bottom). Some plaques (middle right and bottom) also experience large LDL influxes. The plaque at the top is unlikely to become dangerous. Macrophages undergo apoptosis and are ingested and recycled effectively leaving few macrophages with large lipid loads. The two plaques in the middle row are both somewhat pathological. The plaque on the left has inefficient efferocytosis which results in the formation of a necrotic core. The plaque on the right has high rates of LDL influx which creates a plaque where macrophage recruitment is pathologically enhanced and where many macrophages have very high lipid loads. The plaque at the bottom of the diagram is highly pathological with both a high rate of LDL influx and inefficient efferocytosis resulting in a large necrotic core and a large proportion of macrophages carrying heavy lipid loads.

Our model predicts that if lipid export and cell emigration are suppressed [10], then apoptosis and efferocytosis can concentrate monocyte-derived lipid, that comes from endogenous lipid, to extraordinarily large quantities inside the macrophage population via coalescence (Figures 7 and 8). In this way, macrophages with very high lipid loads may be produced independently of LDL. This theory may explain why macrophage foam cells appear in numerous chronic inflammatory conditions such as tuburculosis granulomas [47] as well as in atherosclerotic plaques. Furthermore, spontaneous foam cell formation may produce a positive feedback with monocyte recruitment due to cytokine signalling by foam cells, which sustains a nonresolving inflammatory response [12]. Suppressed lipid export may be beneficial when an inflammatory response is required to clear an infectious agent [10] although we have not included this behaviour in the model. However suppressed lipid export may be detrimental in the to the resolutions of nonresolving inflammatory responses such as to oxidised LDL during atherosclerosis or apoptotic adipocytes in obesity [48]. Efferocytosis can, therefore, act as a double edged sword; it is necessary to prevent postapoptotic necrosis, but may drive excessive lipid accumulation in macrophages.

The inflammatory response inside the artery wall becomes pathogenic when the necrotic core and macrophage population size are large [13]. Our model generates these disease characteristics when the rates of monocyte and LDL influx into the artery wall are large and efferocytosis and emigration are suppressed (see Figure 9).

When efferocytosis is efficient and there is a large influx of LDL, average macrophage lipid loads may become high. This promotes cytokine signalling and so increases the rate of monocyte recruitment and expands both the population size and the amount of lipid in the tissue. Macrophage numbers and lipid contents can be large even while the necrotic core size is small. This is particularly likely when emigration rates are high. In this case efferocytosis clears all apoptotic macrophages and recycles lipid back into the macrophage population and macrophages can ferry their lipid load out of the tissue. This state, where macrophage numbers and lipid content are large, can become vulnerable to becoming pathological if the efferocytosis rate declines. Where the rate of cell emigration is low and efferocytosis is inefficient, a proportion of apoptotic cells will not be ingested by live macrophages, but will become necrotic. The necrotic material will be ingested by macrophages, but if these macrophages can rapidly offload lipid to HDL or emigrate from the plaque, then lipid loads will remain low, despite the presence of the necrotic core. (See Figure 9.)

Cell emigration typically promotes disease regression [18]. Emigration leads to lipid removal from the artery wall as and, thereby, prevents it from being recycled via efferocytosis. Reduction in apoptotic cell formation also reduces the extent of postapoptotic necrosis when efferocytosis is inefficient. In pratice, however, the rate of emigration is small compared to the rates of recruitment and efferocytosis. [8, 49]. A quantitative study suggests that 5 − 15% (Ω = 1.5 − 1.15) of monocyte-derived cells leave via cell emigration during inflammation [8]. Although the proportion of macrophages that undergo emigration is small [8, 49], our model suggests that it is significant in relation to lipid accumulation (Figure 7).

Where a large rate of recruitment is balanced by a large rate of emigration, LDL-derived lipid, apoptotic cells and necrotic debris that have been consumed by macrophages, are effectively moved from the artery wall. High monocyte recruitment may be beneficial as it increases macrophage numbers and so reduces the average lipid load per cell, but recruitment can also be detrimental as it introduces fresh endogenous lipid into the system. Macrophage populations may also grow via cell proliferation [16, 17], although we have not included this in this model. Cell division may be a beneficial way to increase macrophage numbers because it dilutes lipid across the population and acts as a branching or fragmentation process that opposes the reverse-branching or coagulation process associated with efferocytosis. Thus a coagulation-fragmentation process can describe the population dynamics of substance accumulation inside macrophages that divide, die via apoptosis and ingest apoptotic cells via efferocytosis.

Our model can readily be extended to represent changes in macrophage phenotype as a function of lipid load. For example, lipid accumulation increases the rate of macrophage inflammatory signalling, [23], apoptosis [24, 25] and necrosis [26]. This type of behaviour can be easily incorporated allowing the rates of apoptosis, efferocytosis, macrophage emigration and ingestion of LDL in equations (29) to (37) to be functions of lipid load *a*. With these changes, the quantity of cytokines or apoptotic cells in the artery wall, for example, will depend on both the number of macrophages and their distribution across the spectrum of possible lipid loads. In our model we neglect spatial effects although it is clear from observations that macrophage density and lipid load are spatially structured within the artery wall [50]. Introducing another independent variable for space in our model would enable us to study how the macrophage population evolves over time, space and lipid content.

The model presented in this paper includes, for the first time, the distribution of intracellular lipids in the macrophage population of an atherosclerotic plaque and in the associated population of apoptotic macrophages. Tracking intracellular lipids using this model has generated new insight into the importance of endogeneous lipid in foam cell development, the role of emigration and the factors that promote foam cell formation.

## 5 Acknowledgments

We thank Dr. Christina Bursill of the South Australian Health and Medical Research Institute and Dr. Michael Watson of the University of Sydney School of Mathematics and Statistics for helpful discussions. Mary R. Myerscough and Hugh Z. Ford acknowledge support from an Australian Research Council Discovery Project Grant (to Mary R. Myerscough).

